# Macrophages amplify the spontaneous activity of damaged sensory neurons in a human co-culture model of neuropathic pain

**DOI:** 10.1101/2025.06.05.656880

**Authors:** Paschalina Chrysostomidou, Zoe Hore, Domenico Somma, Julia M. Vlachaki Walker, Katy Diallo, Heather F Titterton, Aziza Elmesmari, Mariola Kurowska-Stolarska, Franziska Denk, Greg A Weir

## Abstract

Neuropathic pain is a highly prevalent condition for which treatments are hampered by low efficacy and dose-limiting side-effects. Injury to the somatosensory nervous system causes maladaptive plasticity that initiates and maintains chronic pain. Emerging evidence suggests that inflammatory cells of the innate immune system shape the response of the injured nervous system and thereby contribute to the pathogenesis of pain. Data from preclinical models and human patient biopsies have specifically implicated peripheral macrophage populations for a pro-algesic role, yet how these cell types influence damaged sensory neurons and whether they directly contribute to neuronal hyperexcitability is unclear. Here, we have developed an iPSC co-culture system to study the interactions of macrophages and sensory neurons in a fully humanised experimental model. We found that analogous to endogenous counterparts, iPSC-derived macrophages (iMacs) display a dynamic molecular and functional profile that is highly dependent on neuronal state. Co-culture with injured iPSC-derived sensory neurons (iSNs) induces morphological, gene expression, and secretory profile changes in iMacs that are consistent with the response of macrophages to nerve injury *in vivo*. iMacs in turn amplify spontaneous firing in damaged sensory neurons, implicating macrophages in this cardinal feature of neuropathic pain. These results illustrate the utility of an iPSC-based model to study signalling between these two cell types; they support a role for macrophages in directly amplifying damaged sensory neuron activity and highlight disrupting pathological signalling between these cell types as a promising strategy for future analgesic drug development.

## Introduction

Neuropathic pain affects over 5 million people in the UK^1^, most of whom are failed by current treatments and live with often disabling pain^2^. Our knowledge of the underlying pathophysiology of pain progression has increased rapidly over recent years but this has so far failed to translate to major breakthroughs in treatment approaches. This suggests a need to expand the cell and signalling pathways traditionally targeted and to develop more advanced translational platforms in which new drug targets can be defined and tested.

Pain arises following damage because of maladaptive plasticity within the somatosensory nervous system^3^. Excessive electrical activity of sensory neurons is key to the initiation and maintenance of neuropathic pain^4^, and this has led to direct targeting of these neurons being the primary focus of analgesic development. However, a multitude of interacting cell types occupy the sensory nerve and dorsal root ganglion (DRG, where the cell bodies of sensory neurons are located) and contribute to nervous system plasticity following injury. Within this context, the precise environmental and cellular cues that drive sensory neuron hyperexcitability are not fully understood. Emerging evidence suggests that inflammatory cells of the innate immune system make an important contribution to neuropathic pain, but exactly which cell types are involved, and the critical signalling pathways that they engage are still unclear. Bridging this gap in our knowledge is important as a better understanding may allow us to disrupt pathological signalling for therapeutic gain.

In animal models of peripheral nerve damage, macrophages at the site of injury and in the corresponding DRG increase in number^5–7^ and undergo major changes in gene expression^8–11^. These cells have been implicated in nerve regeneration^12,13^ and depletion studies have suggested that they initiate and maintain neuropathic pain behaviours^6,14^. Macrophage populations have similarly been associated with the human condition, with macrophage-associated genes being found enriched in DRG and nerve trunk tissue from patients with neuropathic pain^15,16^. Using Morton’s neuroma as a model system for human neuropathic pain, a recent study found a positive correlation between greater macrophage presence and higher paroxysmal pain scores in patients and specifically identified an increased intraneural density of CD163^+^MARCO^+^ macrophages in neuroma tissue^7^. However, despite the evidence implying a prominent role for macrophages in neuropathic pain, we do not yet have a good understanding of how macrophages and damaged sensory neurons interact, and crucially, whether macrophages directly contribute to neuronal hyperexcitability.

Our understanding of macrophage involvement in pain pathophysiology has been greatly advanced by the use of experimental animal models. However, most drug targets identified in these models have failed to translate to patient benefit, which could in part be due to differences between the species. This makes a compelling argument to diversify our experimental platforms and to establish relevant testing models based on human tissue. Here, we have developed the first humanised co-culture system of sensory neurons (iSNs) and macrophages (iMacs) derived from induced pluripotent stem cells (iPSCs). Our findings indicate that iSNs shape the morphology and gene expression and secretion of iMacs. This profile is altered to a putatively more pro-inflammatory state upon neuronal injury, and we identify features consistent with the response of macrophages to nerve injury *in vivo*. Finally, we show that iMacs are capable of directly augmenting damaged iSN hyperexcitability, thus replicating a cardinal feature of neuropathic pain. These findings demonstrate that disrupting pathological signalling between macrophages and damaged sensory neurons is a promising approach for future analgesic development. We therefore present our system which could be used as a translationally relevant analgesic discovery and testing platform.

## Methods

### iPSC culture and maintenance

Experiments were conducted with three iPSC lines that have been previously characterised and described; SBAD2 (derived from skin fibroblasts of a 51-year-old male^17^), SBAD3 (derived from skin fibroblasts of a 36-year-old female^18^) and SFCi55ZsG (derived from skin fibroblasts of a female donor^19^). iPSCs were maintained in mTesR Plus media (Stem Cell Technologies, #100-0276) on Matrigel (Corning, #354234) coated dishes. When confluent, cells were passaged as aggregates with ReLeSR (Stem Cell Technologies, #05872). When cells were required to be plated as single cells prior to differentiation, they were instead passaged by Accutase treatment (ThermoFisher, #A1110501), with plating medium supplemented with CEPT cocktail (chroman 1, emricasan, trans-ISRIB; Tocris, #59222C, #M1028 and #I8896 respectively).

### iPSC-derived sensory neuron (iSN) generation

iPSCs were differentiated to iSNs as per^20^, with only minor modifications. Cells were plated on Matrigel treated plates and expanded in mTESR Plus medium until 40-50% confluent, after which they were switched to KSR base medium, which was composed of Knockout-DMEM (ThermoFisher, #10829018), 15% knockout-serum replacement (ThermoFisher, #10828028), 1% Glutamax (ThermoFisher, #35050038) 1% nonessential amino acids (ThermoFisher, #11140050) and 100 μM β-mercaptoethanol (ThermoFisher, #31350-010). This timepoint is denoted as day 0. Neural induction was initiated by adding SMAD inhibitors SB431542 (10 μM, Tocris, #1614) and LDN-193189 (500 nM, Tocris, #8150) until day 5. Three additional small molecules were introduced on day 3; CHIR99021 (3 μM, Tocris, #4423), SU5402 (10 μM, Stratech, #3300) and DAPT (10 μM, Tocris, #2634). The culture medium was gradually transitioned to N2/B27 medium (Neurobasal medium (ThermoFisher, #21103049), 2% B27 supplement (ThermoFisher, #12587010), 1% N2 supplement (ThermoFisher, #17502048), 1% Glutamax) in 25% increments from day 4. Cells were passaged with Accutase and replated onto glass coverslips at day 12 of the differentiation in N2/B27 medium supplemented with four recombinant growth factors at 25ng/ml (BDNF (Peprotech, #450-02), NT3 (Peprotech, #450-03), NGF (Peprotech, #450-01) and GDNF (Peprotech #450-10)). CHIR90221 was included for a further 4 days. Cytosine arabinoside (4 μM, ThermoFisher, #15244774) was added to cells for 18hrs between days 14 and 20 to remove contamination with dividing cells. From day 25 onwards, cells were maintained in N2/B27 Plus medium (Neurobasal Plus medium (ThermoFisher, #A3582901), 2% B27 Plus supplement (Thermofisher, #A3582801), 1% N2 supplement, 1% Glutamax supplemented with 25ng/ml BDNF, GDNF, NT3 and NGF, and Matrigel (1:500). Medium was 50% refreshed 3 times per week and cells used for experiments >45 div. For fluorescent labelling of neurons, AAV.PHP.s.hSyn.mCherry (University of Zurich Viral Vector Facility) was added one week prior to experiments at an MOI of 10^5^.

### iMac generation

iPSCs were differentiated to iMacs as per^18^, with only minor modifications. Single-cell suspensions of 4×10^6^ iPSC cells were seeded into an Aggrewell 800 well (STEMCELL Technologies, #34825) to form embryoid bodies (EBs). These cells were maintained in mTesR Plus media supplemented with 50 ng/mL BMP4 (ThermoFisher, #PHC9534), 50 ng/mL VEGF (ThermoFisher, #PHC9391), and 20 ng/mL SCF (ThermoFisher, #PHC2111). CEPT cocktail was included in the medium on the day of seeding. Media was 75% refreshed daily for 6 days before EBs were plated on Matrigel-treated T25/75 flasks at an approxmate density of one EB/cm^2^. EBs were maintained in flasks in X-VIVO15 media (Lonza, #02-060Q) supplemented with 100ng/ml M-CSF (ThermoFisher, #PHC9501), 25ng/ml IL-3 (ThermoFisher, #PHC0031), 1% Glutamax, 0.055mM β-mercaptoethanol and 1% antibiotic-antimycotic (anti-anti, ThermoFisher, #15240096). Media was 50% refreshed weekly until the emergence of iMac precursors in the supernatant at which point 100% media changes were performed twice weekly. iMac precursors were harvested from the supernatant and plated for 7 days in X-VIVO15 media supplemented with 100ng/ml M-CSF, 1% Glutamax, and 1% anti-anti, to promote final differentiation. To confirm phagocytic activity, pHrodo Red S. aureus BioParticles (ThermoFisher, #P35367) were added to iMacs at 50 μg/ml and fluorescence imaged every 30mins on a Zoe Fluorescent Cell Imager (BioRad).

### Monocyte-derived macrophages (MoMs)

30ml of peripheral blood was collected from a healthy 47yr-old female, in accordance with the University of Glasgow’s MVLS Ethics Committee (Project No: 2012073), and with written consent. PBMCs were isolated following density gradient centrifugation with Histopaque-1077 (Sigma Aldrich, #10771) and CD14^Pos^ cells were isolated using CD14 microbeads and AutoMACSPro (Miltenyi BioTec, #130-050-201), according to the manufacturer’s protocol. These cells were plated into six-well culture plates and differentiated into monocyte-derived macrophages by 7 days of culture in RPMI 1640 medium (ThermoFisher, #31870-025) containing 10% foetal bovine serum (ThermoFisher, #10439001), 1% anti-anti and 50ng/ml M-CSF (complete RPMI media). Media was replenished on day 3 and cells increased their adherence throughout the 7 days.

### Co-culture of iSNs and iMacs/MoMs

iMacs/MoMs were added to iSNs in a 1:1 ratio in N2/B27 Plus medium (Neurobasal Plus medium, 2% B27 Plus supplement, 1% N2 supplement, 1% Glutamax) supplemented with 25ng/ml BDNF, GDNF, NT3 and NGF, and 10ng/ml M-CSF. Co-cultures were initiated when iSNs were mature (>45 div) and in the case of our injury model, 18hrs after iSN axotomy. 50% media changes were performed twice per week, and co-cultures were maintained for 1-42 days.

### iSN axotomy

Axotomy of iSNs was performed similarly to previously described^21^. Mature iSNs were manually lifted from their growth plates and dissected into smaller aggregates with a scalpel blade. Cell clusters were then mechanically dissociated to single cells by using fire-polished glass pipettes of decreasing bore size. The subsequent cell suspension was then replated onto Matrigel coated glass and allowed to adhere for 18hrs before addition of iMacs.

### Sample preparation for single cell RNA sequencing of iMacs/MoMs

Cells from all conditions were generated, collected, processed and sequenced in parallel. On the day of isolation, cultures were enzymatically dissociated with 30 minutes of Accutase (ThermoFisher) followed by brief mechanical dissociation with a fire polished pipette. Single cell suspensions were transferred to phosphate buffered saline (PBS) containing 1:1000 eBioscience™ Fixable Viability Dye eFluor™ 780 (ThermoFisher, #65-0865-14) and incubated on ice for 20 minutes. Samples were maintained in dark conditions from this point onwards. Samples were centrifuged and washed in fluorescence-activated cell sorting (FACS) buffer containing PBS, 2% foetal bovine-serum, 1% anti-anti and 0.5mM EDTA (ThermoFisher, #15825388) before being resuspended in Human TrueStain FcX blocking buffer (BioLegend, #422301) diluted 1:10 in FACS buffer and incubated for 10 minutes on ice. Brilliant Violet 711 anti-human CD45 (BioLegend, #304049) (1:100) and a unique TotalSeq-A hashtag oligo (HTO) (A-0251 (BioLegend, #394601), A-0252 (BioLegend, #394603), A-0253 (BioLegend, #394605), A-0255 (BioLegend, #394609), A-0256 (BioLegend, #394611), A-0258 (BioLegend, #394615), A-0259 (BioLegend, #394617), A-0260 (BioLegend, #394619)) were added directly to each tube and incubated for 30 minutes on ice. Samples were subsequently centrifuged, resuspended in FACS buffer and strained (100µm strainer) into FACS tubes. Sorting was performed by an BD FACSAria III within the University of Glasgow Cellular Analysis Facility. Gating strategy is documented in Supp Fig 4. Unstained cells, Viability Dye eFluor™ 780 stained cells and single stained control beads were used for compensation. Cells were sorted into 1.5ml microcentrifuge tubes containing complete RPMI media (see MoMs section).

### Single cell RNA sequencing (scRNA-seq) of iMacs/MoMs

Sorted cells were manually counted with a haemocytometer and 2,500 cells per condition were mixed for a total of 20,000 cells. Cells were loaded onto a Chromium Controller (10X Genomics) for single-cell partitioning, using Chromium Next GEM Chip G Single Cell Kit (10X Genomics #PN-1000120), and single-cell 3’ Reagent Kits v3.1 (10X Genomics #PN-1000121). A library consisting of iMacs and MoMs was sequenced on an Illumina Nextseq 500 according to the manufacturer’s protocol, at the Glasgow Polyomics facility at a depth of 20,000 reads/cell. Reads were mapped to the human genome (GRCh38-3.0.0) using 10X CellRanger (version 7.0.0).

### scRNA-seq analysis of iMacs/MoMs

Analysis was carried out using the Seurat package (version 5.2.1) in RStudio (version 4.4.2). Briefly, the data were loaded using the Read10X() and CreateSeuratObject() functions. Prior to downstream analysis, the library was filtered to only retain cells with > 200 and < 5000 unique genes as well as cells in which < 10% of the counts were from mitochondrial genes for downstream analysis. Counts were subsequently scaled to 10,000 transcripts per cell using the NormalizeData() function and highly variable genes were identified using FindVariableFeatures(). The data were then scaled to equalise variance amongst genes and dimensionality reduction was performed using the RunPCA() function. A shared K-nearest neighbour (KNN) graph was constructed to identify the nearest neighbours for each cell based on the first 20 principal components (PCs) using the FindNeighbours() function and clustering was performed using FindCusters() at a resolution of 0.2. For visualisation, uniform manifold approximation and projection (UMAP) were generated with coordinates calculated from the first 20 PCs using the RunUMAP() function.

Prior to further analysis, samples were deconvoluted by the addition of HTO information using cellhashR (v 1.0.3) (Table 1). Cells with no hashtag information and doublets were removed, and the UMAP was re-generated using the pipeline described above. The library was further divided into two distinct datasets, iMacs and MoMs, each containing macrophages from four conditions. These were analysed separately following the steps described previously. Clustering was performed at a resolution for 0.1 for the iMac and 0.15 for the MoM datasets. Differential expression analysis was carried out using the FindAllMarkers() function (min.pct=0.25, test=“MAST”) and clusters were labelled based on the top 15 differentially expressed genes (FDR < 0.05). All raw and processed sequencing files, including Seurat objects, can be obtained from the Gene Expression Omnibus (GEO) under accession GSE298325. The full analysis code is provided in Supplementary RNotebook 1.

**Table 1.**
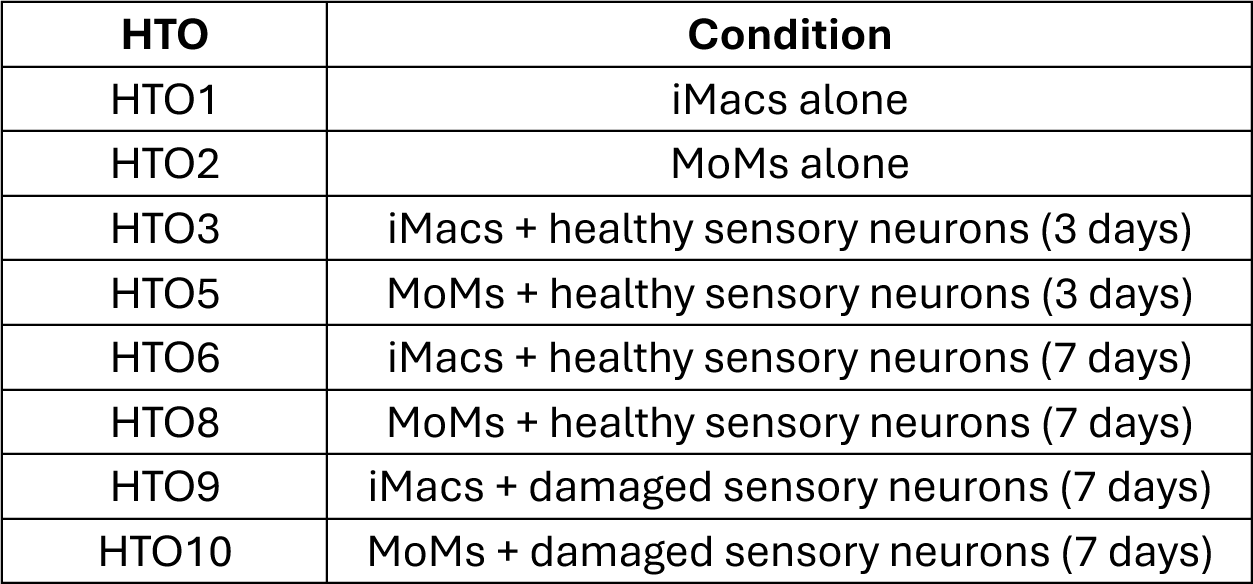
Hashtag oligo (HTO) barcodes used in CITE-seq multiplexing and their corresponding conditions.

### Cytokine/chemokine release assays

A custom U-PLEX assay platform (Meso Scale Discovery) was used as per manufacturer’s instructions to profile the abundance of fourteen cytokines/chemokines in the supernatant of cultures containing either iMacs only, iMacs with healthy iSNs, iMacs with damaged iSNs, healthy iSNs only, or damaged iSNs only. Supernatant was taken after 3 days of culture and centrifuged to remove debris, before profiling using the U-Plex assay. Plates were read by a MESO SECTOR S 600 Reader. A standard curve was derived from signals from standards and generated with a four-parameter logistic model. Sample concentrations were determined by backfitting to the standard curve and multiplied by the dilution factor. Galectin-3 (ThermoFisher, #BMS279-4), CCL3 (ThermoFisher, #88-7035-22), and CCL4 (ThermoFisher, #88-7034-88) were profiled using human ELISA kits according to manufacturer’s instructions. Signal intensity was determined using a Multiskan Spectrum plate reader (ThermoFisher) and sample concentration determined using a standard curve.

### Immunocytochemistry

iMacs and iSNs, both co-cultured and alone, were fixed either directly with 4% PFA for 15 minutes at room temperature, or by overlaying culture media with 4% PFA containing 10% sucrose for 1 hour at room temperature. Coverslips were washed three times with PBS before incubating with primary antibodies diluted in PBS with 0.3% Triton™ X-100 (ThermoFisher, #A16046.AE) and 10% normal donkey serum for 2 hours at room temperature. After 3 PBS washes, cells were incubated with secondary antibodies for 2 hours at room temperature before being stained with DAPI (Sigma, #D9542) for 10 mins. Coverslips were then washed 3 times with PBS, mounted onto Vectashield antifade medium (Vector Laboratories, #H-1400-10) and allowed to harden for 1 hour at room temperature before imaging. Primary antibodies used in this study: α-NF200 (1:1000, mouse, Sigma-Aldrich #266004), α-BRN3A (1:100, rabbit, Merck #AB5945), IBA1 (1:500, guinea pig, Synaptic systems #234 308), α-GFP (1:1000, chicken, Abcam #ab13970), α-ATF3 (1:500, rabbit, Novus Biologicals #NBP1-85816), α-FABP5 (1:200, rabbit, ProteinTech #12348-1-AP), and α-β-tubulin III (1:1000, mouse, Sigma-Aldrich #N5389). A range of species-specific secondary antibodies were used that were conjugated to Pacific Blue, Alexa fluor 488, Rhodamine Red-X, Alexa fluor 647 (Jackson ImmunoResearch) and Alexa fluor 488+ (ThermoScientific).

### Image acquisition and analysis

All fluorescent images were acquired with a Zeiss LSM900 Airyscan confocal microscope equipped with 405-, 488-, 561-, and 640-nm diode lasers and analyses were performed using FIJI. *Quantification of ATF3 signal*: iMac nuclei from cells cultured with healthy or damaged iSNs were identified by IBA1 and DAPI and labelled for region of interest (ROI) analysis, blind to ATF3 signal. Nuclei masks were then overlayed onto the ATF3 channel, and the mean grey area of each ROI was automatically calculated in FIJI. Values were aggregated across cells from multiple fields of view to derive an average ATF3 signal intensity per sample. *Quantification of FABP5 signal*: iMac cell bodies from cells cultured with healthy or damaged iSNs were identified by IBA1 and labelled for ROI analysis, blind to FABP5 signal. Cell masks were then overlayed onto the FABP5 channel, and the mean grey area of each ROI was automatically calculated in FIJI. Values were aggregated across cells from multiple fields of view to derive an average FABP5 signal intensity per sample. To mitigate confounds of batch processing for both ATF3 and FABP5 quantification, culture, fixation, ICC and image acquisition were run in parallel for each group from a given differentiation. For iMac morphology, a cell was classified as positive for secondary branching if any processes were observed extending from primary processes. For cell size analysis, IBA1 positive cells were outlined using FIJI and identified using the processing steps: *threshold-despeckle - make binary-convert to mask - fill holes - wand selection*. Any cell which was not accurately outlined using this automated procedure was manually traced. ROIs were subsequently quantified for their area.

### Whole-cell patch clamp electrophysiology

Whole-cell patch clamp recordings of iSN were performed at room temperature (22°C) with a Multiclamp 700B amplifier and Digidata 1550 acquisition system (Molecular Devices). Data were low pass filtered at 2 kHz and sampled at 10 kHz and recordings were analysed by Clampfit 10 software (Molecular Devices). Filamental borosilicate glass capillaries (1.5 mm OD, 0.84 mm ID; World Precision Instruments) were pulled to form patch pipettes of 3–7 MΩ tip resistance and filled with internal solution containing (mM): 130 K-gluconate, 10 KCl, 2 MgCl_2_, 2 ATP-Na_2_, 0.5 GTP-Na, 10 HEPES, and 0.5 EGTA; pH was adjusted to 7.3 with KOH and osmolarity set to 305 mOsm. Extracellular solution contained (mM): 140 NaCl, 3 KCl, 2 MgCl_2_, 2 CaCl_2_, 10 HEPES and 10 glucose; pH was adjusted to 7.3 with NaOH and osmolarity was set to 315 mOsm. Resting membrane potential values were corrected offline for a liquid junction potential of −15.5mV. One minute following establishing whole cell access, a two-minute recording was made in bridge mode to derive resting membrane potential and spontaneous action potential firing. Input resistance (R_Input_) was derived from the membrane potential deflection caused by a 20pA hyperpolarising current pulse at −60 mV. To determine the minimal current injection to fire an action potential (rheobase), cells were depolarised from a holding potential of −60 mV by current steps (50 ms) of increasing magnitude (Δ10 pA) until an action potential was generated. Recordings were discarded in cases of poor whole-cell breakthrough (evident from excessive leak current or recording instability), or if neurons failed to fire an action potential to any level of current.

### *In vivo* scRNA-seq dataset of mouse myeloid cells

#### Surgical nerve damage in mice

Male and female C57BL/6JolaHsd mice were purchased from Envigo at 7 weeks of age and housed in groups of up to five in individually ventilated cages in a 12h light-dark cycle. All experiments were carried out in accordance with the United Kingdom Home Office Legislation (Scientific Procedures Act, 1986) and were approved by the Home Office to be carried out at King’s College London. Mice underwent one of two models of nerve damage: partial sciatic nerve ligation (PSNL) or sciatic nerve crush. For both procedures, animals were anaesthetised under 2% isoflurane and pre-operatively administered 0.1mg/kg of the analgesic buprenorphine subcutaneously (s.c). After shaving the fur covering the left hip and leg region, the sciatic nerve was exposed via blunt dissection. For PSNL, roughly half of the nerve was tightly ligated with a suture (5.0 Vicryl suture, Ethicon, #W9982). For nerve crush, Watchmaker forceps (Dumont, #0304-5-PO) were placed horizontally around the nerve and pressed tightly for 10 seconds, the forceps were then placed vertically around the same section of nerve and again pressed tightly for 10 seconds. For sham surgeries the sciatic nerve was exposed but left untouched. In all cases, the wound was stapled shut with a Reflex 7mm metal wound clip (Fine Science Tools, #12032-07). Mice were transferred to a clean cage and left to recover in the animal unit. Animals were weighed and checked for autotomy daily for at least 3 days following surgery and wound clips were removed within 7 days of surgery.

#### Nerve dissociation and processing

1 week or 15 weeks after surgery, mice were transcardially perfused with 10 ml PBS to remove leukocytes from circulating blood. Ligated, crushed or uninjured (sham) nerves were dissected and placed into a petri dish containing F12 medium (Gibco, #21765-029). Ligatures were removed with tweezers, if still present, and nerves were cut to 0.5 cm, ensuring equal lengths of nerve either side of the ligation/crush site. Following dissections, samples were transferred into wells of a standard 96-well PCR plate containing 51μl of the following digestion mix: 6.25 mg/ml collagenase type IA (Sigma Aldrich, #C9891), 0.2% Pronase (Millipore, #53702), 0.4% Hyaluronidase (ABNOVA, #P52330) in F12 medium. Spring scissors were used to chop the tissue into small pieces (50x chops, cleaning scissors between samples with 100% ethanol). After this manual dissociation, the plate was incubated on a shaking platform (220 rotations per minute) at 37°C for 30 minutes. The digestion mix was subsequently removed and replaced with 100μl of fluorescence-activated cell sorting (FACS) buffer: 0.4% BSA (Sigma-Aldrich, #A3983), 15 mM HEPES (Gibco, #15630080), 2mM EDTA (Invitrogen, #15575038) in Hank’s Buffered Saline Solution (HBSS, diluted from 10x, Gibco, #14175095). Samples were then further dissociated by pipetting up and down 50x, with the resulting single-cell solution filtered through a 35μm cap of a BD falcon tube with a cell strainer cap (BDBiosciences, # 352235). The plate was centrifuged at 1500 rotations per minute for 5 minutes at 4°C, after which supernatants were removed and resuspended in 50μl of the following staining mixture for 30 minutes at 4°C in a dark fridge: Fc block (BioLegend, #101302, 1:200), FITC-CD45 to stain leukocytes (BioLegend, #147709, 1:1200), PE-Cy7-CD11b to stain myeloid lineage cells (BioLegend, #101215, 1:1200) and, depending on the sample, one of four hashtag antibodies: TotalSeq-B0301-MHCI (BioLegend, #155831), TotalSeq-B0302-MHCI (BioLegend, #155833), TotalSeq-B0303-MHCI (BioLegend, #155835), TotalSeq-B0304-MHCI (BioLegend, #155837). At the acute time point, one week after surgery, the antibody staining mix also contained BUV395-Ly6G to stain neutrophils (BD Bioscience, #563978, 1:300). After staining, cells were centrifuged for 5 minutes at 1,500 rotations per minute at 4°C, after which the cell pellets were resuspended in 300μl FACS buffer and transferred to FACS tubes, adding DAPI to stain for dead cells (Invitrogen, #D1306, 1μl of a 1:2 dilution).

#### FACS of mouse nerves

Sorting was performed on a BD FACSAria II or BD FACSAria Fusion in the Advanced Cytometry Platform Core Facility of the Guy’s and St Thomas’ NHS Foundation Trust. Gating strategies are shown in Supp figs 6 and 7. Unstained cells, DAPI single stained cells and single stained control beads (BD™ CompBeads, BD Bioscience, #552845) were used for compensation. Cells were sorted into 1.5ml microcentrifuge tubes containing 4μl of 4% BSA (Sigma-Aldrich, #A3983) in PBS. All tubes were pre-coated for at least 1 hour with the same 4% BSA solution and placed on ice immediately following cell collection. The following samples were collected: for the 1-week time point, CD45+/CD11b+/Ly6G-live single cells from n = 2 sham (2,000 cells each), crush and PSNL mice (8,000 cells each); for the 15-week time point, as many CD45+/CD11b+ live single cells as possible were collected from n = 3 crush and n = 4 PSNL mice.

#### scRNAseq of mouse myeloid cells

After FACS, cells were immediately taken to the Guy’s and St Thomas’ Genomics Core Facility which was located on the same floor as the Advanced Cytometry Platform. Hashed samples were combined as follows in two separate 10X runs: for the 1-week time point, one 10X well (acute crush) was filled with cells from 1x male sham, 1x male crush, 1x female crush mouse, while the other (acute PSNL) was filled with cells from 1x female sham, 1 male PSNL and 1 female PSNL mouse. A separate 10X run was performed for the 15-week time point, with one well (chronic crush) filled with 2x female crush and 1x male crush sample, and the other well (chronic PSNL) filled with 2x female and 2x male PSNL samples. Libraries were generated using the 10X Genomics Chromium Single Cell 3ʹ Feature Barcode Library & Gel Bead Kit (v3 Chemistry) according to manufacturer’s instructions and sequenced on an Illumina NextSeq 2000 P2 Flow Cell obtaining mean reads per cell of: 24,752 (acute crush well), 35,710 (acute PSNL well), 45,531 (chronic crush well), and 59,220 (chronic PSNL well).

#### Data analysis of mouse scRNA-seq

Reads were aligned using the 10X Genomics Cell Ranger (version 6.0.0) “cellranger count” pipeline, using the pre-built mouse GRCm38 transcriptome reference and following guidance for “Feature Barcode Analysis” using TotalSeq-B antibodies. The following cell numbers were captured: 1,920 (acute crush), 2,471 (acute PSNL), 2,675 (chronic crush), 2,067 (chronic PSNL). Output files from Cell Ranger were analysed in R Studio (Version 2024.09.1) with Seurat (version 5.1.0)^22^. Within each time point, samples were run on the same 10X chip and sequenced in parallel, minimizing technical batch effects. We therefore first analysed the acute and chronic timepoint data separately. The following cells were excluded to eliminate any empty cells or potential duplets: cells with less than 150 or more than 4,000 (acute) or 4,500 (chronic) unique molecular identifiers (UMIs); cells with more than 20,000 molecules (acute only); cells with more than 10% mitochondrial gene content. Cells negative for a hashtag or positive for more than a single hashtag were also excluded. Cluster identity was assigned based on known marker gene expression. To combine acute and chronic timepoints, three different integration methods were compared in Seurat, with reciprocal PCA deemed to best replicate the clusters that emerged from the individual time-point analyses. See Supplementary RNotebook 2 and its corresponding html file for analysis code and output. For comparison with the human macrophage scRNAseq data, we used the UCell package^23^ to project marker genes for the SPP1^high^ and SPP1^low^ iMac clusters onto the mouse scRNAseq UMAP space. Marker genes were identified using the Seurat FindAllMarker function; to be included in the analysis, they had to expressed in at least 10% of cells in each cluster and had to differ in their expression from the other clusters by at least 20% (SPP1^low^ MRC1^pos^ and mixed SPP1^high^ MRC1^pos^ clusters) or 30% (SPP1^high^ MRC1^neg^, cycling cell clusters) at adjusted p < 0.05. See Supplementary Data Table 4 for the gene lists we obtained and Supplementary RNotebook 2 for the analysis code. We also made a Sankey plot to directly compare mouse and human scRNA-seq clusters. For this, we identified human orthologues of the mouse genes in our data (Supplementary RNotebook 2). Finally, we performed ligand receptor analysis using the ICELLNET package^24^, investigating which ligands on our human iMacs would be able to bind receptors present on iSNs. For this, we used previously published RNA sequencing data of iSNs differentiated for 30, 50 and 70 days^25^. Only genes that were deemed expressed on iSNs in this dataset, as well as genes that were expressed in at least 10% of iMac clusters were included in the analysis. See Supplementary RNotebook 3 and its html output for the script that was used. All raw and processed sequencing files, including Seurat objects, can be obtained from the Gene Expression Omnibus (GEO) under accession number GSE298326.

### Quantification and statistical analysis

Data are expressed throughout as mean ± standard error of the mean (SEM). Statistical testing was performed with GraphPad Prism with significance for all experiments placed at p < 0.05. A Student’s *t*-test was used to compare the mean of two groups and when data were not normally distributed a non-parametric test was applied (Mann-Whitney). Electrophysiological measures were assessed by two two-way ANOVA analysis, testing for main effects of injury and iMac presence, and subsequent Tukey’s post-hoc tests for pairwise comparisons. Sample sizes are detailed in each figure legend.

## Results

### Establishing a humanised co-culture model

With the goal of generating co-cultures of human sensory neurons and macrophages, we took advantage of previously characterised iPSC differentiation protocols. Human iPSCs can be differentiated into neurons with a transcriptional and functional profile consistent with those of human DRG neurons (iSNs) by using a cocktail of small molecule inhibitors (Fig. 1a)^20,26^. The precise sensory neuron subtype that they most closely resemble is unclear; however, they exhibit features of nociceptors^17,20^ and have proved a valuable model to investigate the physiology of human sensory neurons^17,27–29^. We generated iSNs from one healthy control iPSC line (SBAD2)^17^. These robustly differentiated into a homogeneous culture of neurons with extensive neurite outgrowth, and expressed the sensory neuron marker, BRN3A (Fig. 1b). Concomitantly, we used a well-characterised embryoid body-based protocol to generate primitive macrophages from iPSCs (iMacs) in a process reminiscent of yolk sac haematopoiesis^18,30^. iMacs were generated from three healthy control iPSC lines (SBAD2, SBAD3^17^ and a previously characterised ZsGreen knockin-line^19^). We confirmed successful iMac differentiation for all three lines based on morphology, expression of canonical genes such as IBA1, CD45 and CD64, and phagocytic competence (Fig. 1c and Supp. Fig.1a-e). Macrophages are highly plastic, and their profile is dependent on the tissue in which they reside. iMacs are similarly able to acquire specific tissue-resident macrophage phenotypes in co-culture environments^18,31^ but until now they have not been co-cultured with sensory neurons. iSNs and iMacs co-cultured in neuronal-based media with the addition of low levels of Macrophage Colony Stimulating Factor (M-CSF) were viable, and iMacs largely associated with clusters of iSN cell bodies (Fig. 1d). While iMacs cultured alone in neuronal media exhibited an elongated or amoeboid morphology (Fig. 1e), iMacs associated with iSN cell bodies took on a more complex morphology, including a reduced size with an increased number of processes (Fig. 1f-h), consistent with the ramified morphology of endoneurial macrophages in the sciatic nerve^32,33^. Co-culture successfully maintained the iMac populations for long periods (6 weeks), with iMac morphology appearing to increase in complexity at these time points (Fig. 1i). These data demonstrate the feasibility of co-culturing these two cell types and suggest a functional interaction between them that alters the iMac state.

**Figure 1.**
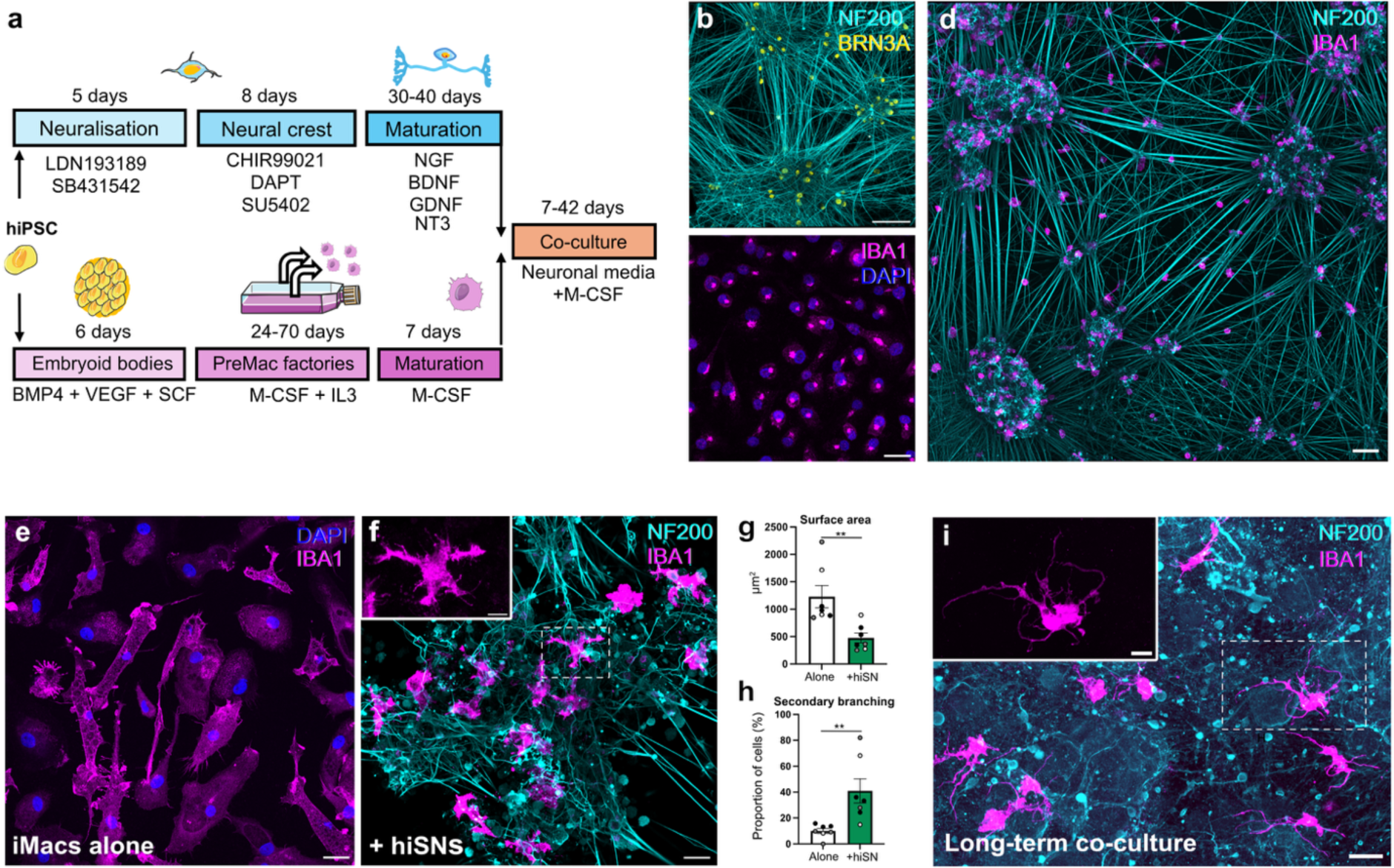
Co-cultures are viable and induce a more ramified iMac morphology. **a** Schematic overview of the experimental setup in which iPSCs are differentiated in parallel to sensory neuron-like cells (iSNs)^20^ and macrophage-like cells (iMacs)^30^, before co-culture in defined media (see *Methods*). **b** 50 days in vitro (div) iSNs expressing the sensory neuron marker, BRN3A, and exhibiting extensive neurite growth (NF200). *Scale bar represents 100*µm. **c** iMac precursors expressing the myeloid cell marker, IBA1. **d** Representative image of iSNs and iMacs co-cultured for 14 days, illustrating the tendency of iMacs to associate with groups of iSN cell bodies. *Scale bar represents 100*µm. **e** Representative morphology of iMacs cultured for 7 days in the absence or **f** presence of healthy iSNs (hiSNs). **g, h** Co-culture reduces iMac cell size and results in a more ramified morphology. Data represents mean ± SEM, with values for individual differentiations also shown (n=7, from 3 independent iPSC lines: *black circles - SBAD2, grey circles - SBAD3 and white circles - ZsGreen*). Mann-Whitney test (**g**) and two-tailed unpaired *t*-test (**h**). \*\**P*<0.01. **i** Representative image of cells in co-culture for 42 days, showing that the two cell types co-exist for long-periods and iMacs retain a ramified morphology. *Inset scale bars represent 10*µm and *all other scale bars represent 25µm unless specified otherwise*.

### Neuron damage alters iMac morphology and secretory profile

Following peripheral nerve damage *in vivo*, nerve and DRG-associated macrophages, and microglia in the dorsal horn undergo major changes in morphology and gene expression^8,10,34^. These changes are thought to be driven, at least in part, by soluble factors released directly by damaged sensory neurons, such as CSF1^34,35^. To study how neuronal damage altered the interaction between our cells *in vitro*, we employed a mechanical trauma nerve injury model^21^. Here, mature iSNs are dissociated and replated, resulting in complete axotomy (Fig. 2a). Damaged iSNs robustly express the canonical injury gene, ATF3, and rapidly regenerate neurites (Supp. Fig. 2a), mirroring the regenerative capacity of the peripheral nervous system. *In vivo*, resident macrophages within the sciatic nerve typically enlarge, retract their processes, and take on a more rounded morphology following nerve damage^32,33,36^. iMacs co-cultured with damaged iSNs for 7 days formed dense clusters around neuronal cell bodies (Fig. 2b). These clusters had reduced adherence, making the standard processing for immunocytochemistry (ICC) challenging. We therefore repeated experiments in co-cultures of iSNs transduced with AAV9-hSyn-mCherry and ZsGreen iMacs, which allowed us to monitor cell dynamics by endogenous fluorescence (Supp. Fig. 2b and c). Most iMacs were observed in neuron soma-associated clusters and appeared to exhibit a more ameboid profile (Fig. 2b and Supp Fig. 2ci), although quantification of individual cell morphology was hindered by the density of clusters. iMacs not associated with clusters remained amongst the neurites and typically were large with a flattened morphology (Supp. Fig. 2cii). Our axotomy model resulted in a large amount of cellular debris, composed of severed neurites and dead neurons. Addition of iMacs to cultures of damaged iSNs starkly reduced the amount of this debris (Fig. 2c), presumably via phagocytosis. To directly test this, we labelled iSNs prior to axotomy with AAV-hSyn-eGFP and could observe GFP^+^ particles within iMacs as early as 24hrs after co-culture, suggesting that iMacs actively phagocytosed neuronal debris from the cultures (Fig. 2d).

**Figure 2.**
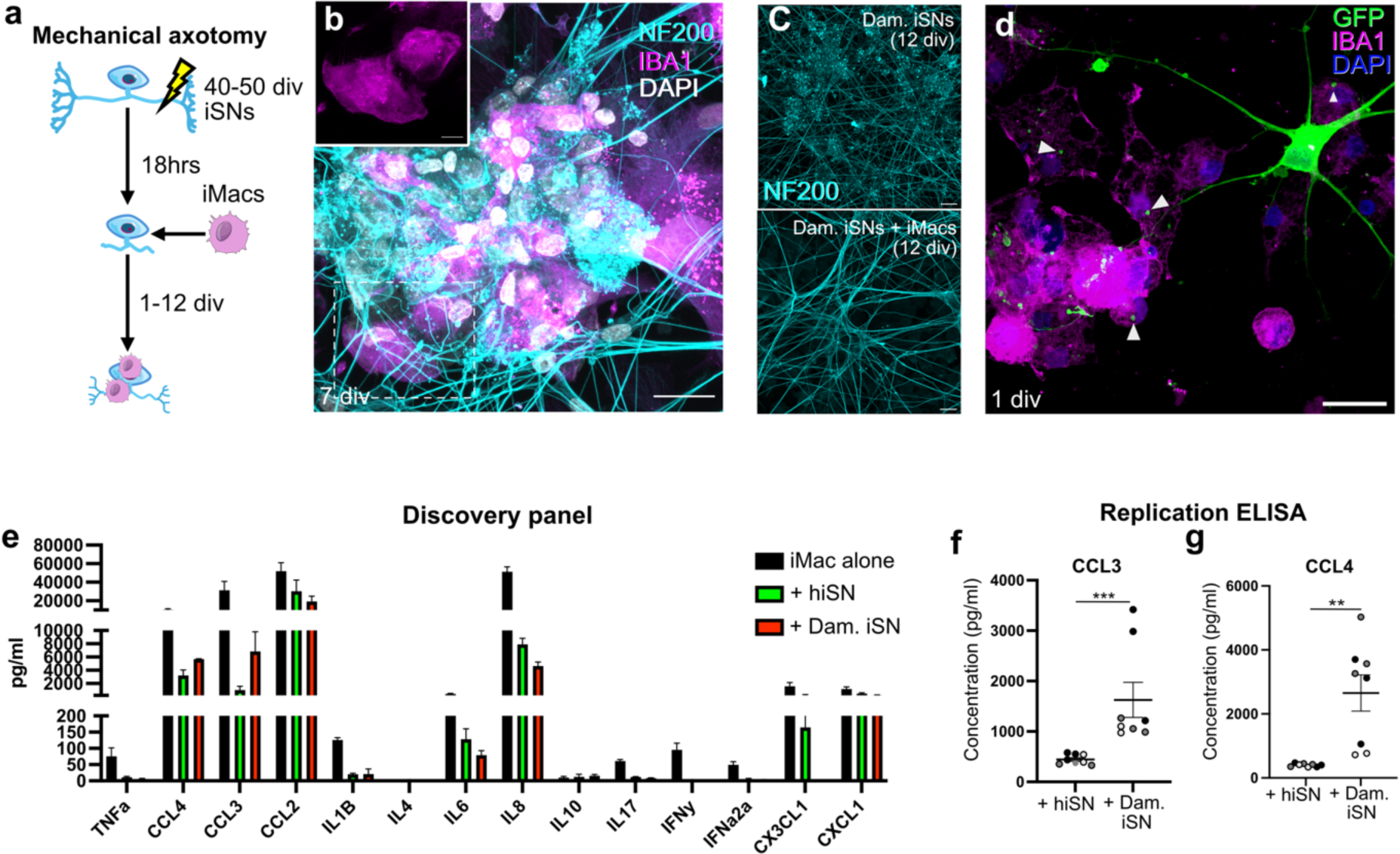
Mechanical injury to iSNs alters iMac morphology and secretory profile. **a** Schematic of the mechanical trauma model in which mature (div 40-50) iSNs are mechanically dissociated and replated, resulting in axotomy. In co-culture experiments, iMacs are added to regenerating iSNs 18hrs later. **b** Representative image of iMacs forming dense clusters with damaged iSN (Dam. iSN) cell bodies following 7 days of co-culture. *Scale bar represents 30*µm *in main image and 10 µm in inset.* **c** iSNs 12 days after damage cultured with and without iMacs. The addition of iMacs considerably reduces the amount of debris in damaged iSN cultures. *Scale bars represent 25*µm. **d** Image of a damaged iSN transduced with AAV-hSyn-eGFP and cultured with iMacs. Arrows highlight GFP^+^ particles within iMacs one day after co-culture, indicating active phagocytosis of neuronal debris. *Scale bar represents 25*µm. **e** Discovery multiplex immunoassay detecting pain-relevant cytokines/chemokines in the supernatant 3 days following co-culture. Data represent mean ± SEM (n=3, from SBAD2 and SBAD3 iPSC lines). **f,g** Individual CCL3 and CCL4 profiling in a verification cohort of supernatant samples taken following 3 days of co-culture. Data represent mean ± SEM, with values for individual differentiations also shown (n=8, from 3 independent iPSC lines: *black circles - SBAD2, grey circles - SBAD3 and white circles - ZsGreen*). Two-tailed unpaired *t*-test (CCL4) and Mann-Whitney test (CCL3). \*\**P*<0.01, \*\*\**P*<0.001.

Next, we asked whether the secretory profile of iMacs was also neuron state dependent by performing a discovery (n=3) multiplex immunoassay for 14 soluble factors with previous association to nerve injury and pain. Appreciable levels of several soluble mediators were detected in iMac-containing cultures but not in monocultures of either healthy or damaged iSNs (Fig. 2e and Supp. Fig. 3), suggesting that the source of secretion was iMacs. Similar to previous reports^18^, we found that iMacs cultured alone generally secreted more of the cytokines and chemokines than iMacs co-cultured with healthy or damaged iSNs (Fig. 2e). Within our targeted panel, iMacs secreted more CCL3 and CCL4 when co-cultured with damaged compared to healthy iSNs, which we verified in single-target immunoassays performed on independent and larger sample sizes (Fig. 2f and g). Both chemokines contribute to neuropathic pain behaviours in mouse models of nerve injury^37,38^. Taken together, these data suggest that co-culture with damaged iSNs drives a morphological and secretory iMac profile that is distinct from that directed by healthy iSNs.

### Neuron state shapes iMac transcriptional profiles

Macrophage gene expression is highly plastic and dependent on the local tissue environment and context (e.g. injury *vs* no injury). We performed a hypothesis-generating, single-replicate experiment to explore the global gene expression profile of iMacs cultured under different conditions. SBAD2 iMacs were cultured alone, with healthy iSNs for 3 or 7 days, or cultured with damaged iSNs for 7 days. iMac samples were subsequently purified from co-cultures by FACS based on viability and CD45 expression (Supp Fig. 4). Cells from each condition were labelled with an identifying hashtag antibody before being pooled into a single tube (to reduce sequencing variability) and scRNA-seq was then performed. Unbiased clustering separated iMacs into four discrete clusters marked by varying levels of SPP1, which encodes osteopontin, MRC1, which encodes the mannose receptor CD206, and MKI67, which marks actively cycling cells (Fig. 3a-b). Of note, SPP1 is up-regulated in several disease-associated macrophage/microglial states, including central nervous system (CNS) degeneration, ageing and injury^39,40^ and MRC1 expression has been used to identify tissue resident macrophages implicated in the resolution of pain following inflammation and surgical wound injury^41,42^. Demultiplexing of the cells based on hashtag identifier revealed that molecular clusters were broadly represented by discrete culture conditions. iMacs cultured alone were classified as SPP1^Low^MRC1^pos^, while iMacs cultured with healthy iSNs were represented in SPP1^High^MRC1^pos^ cycling and non-cycling populations. iMacs cultured with damaged iSNs were found to be SPP1^high^MRC1^neg^ (Fig. 3c). In addition to these marker genes, cells from each condition could be defined by distinct patterns of gene expression (Fig. 3d). We explored whether iMac gene expression was dependent on iSN state (healthy *vs* damaged) and performed hypothesis-generating (owing to the single biological replicate) pseudo-bulk analysis to compare the two conditions. iMacs cultured with damaged iSNs were enriched for expression of several genes that characterise a neurodegenerative disease-associated microglial (DAM) state (such as ABCA1, LPL, CD9 and APOC1^43^) and those associated with pro-inflammatory conditions (such as C15orf48, CXCL8, NR1H3 and SLAMF7), while they exhibited a concomitant downregulation of genes associated with alternatively activated macrophage populations (such as DAB2, STAB1 and MMP12^44–46^) (Supplementary data table 1). Several of the genes enriched in iMacs co-cultured with damaged iSNs have previously been implicated in neuropathic pain. We chose to validate increased expression of three of these genes that are known to play a role in macrophage biology; LGAL3 (encoding galectin-3), ATF3 and FABP5^47–52^. iMacs secreted greater levels of galectin-3 and exhibited higher protein expression of ATF3 and FABP5 when co-cultured with damaged vs healthy iSNs, across biological replicates (SBAD2, SBAD3 and ZsGreen derived) (Fig. 3e-g).

**Figure 3.**
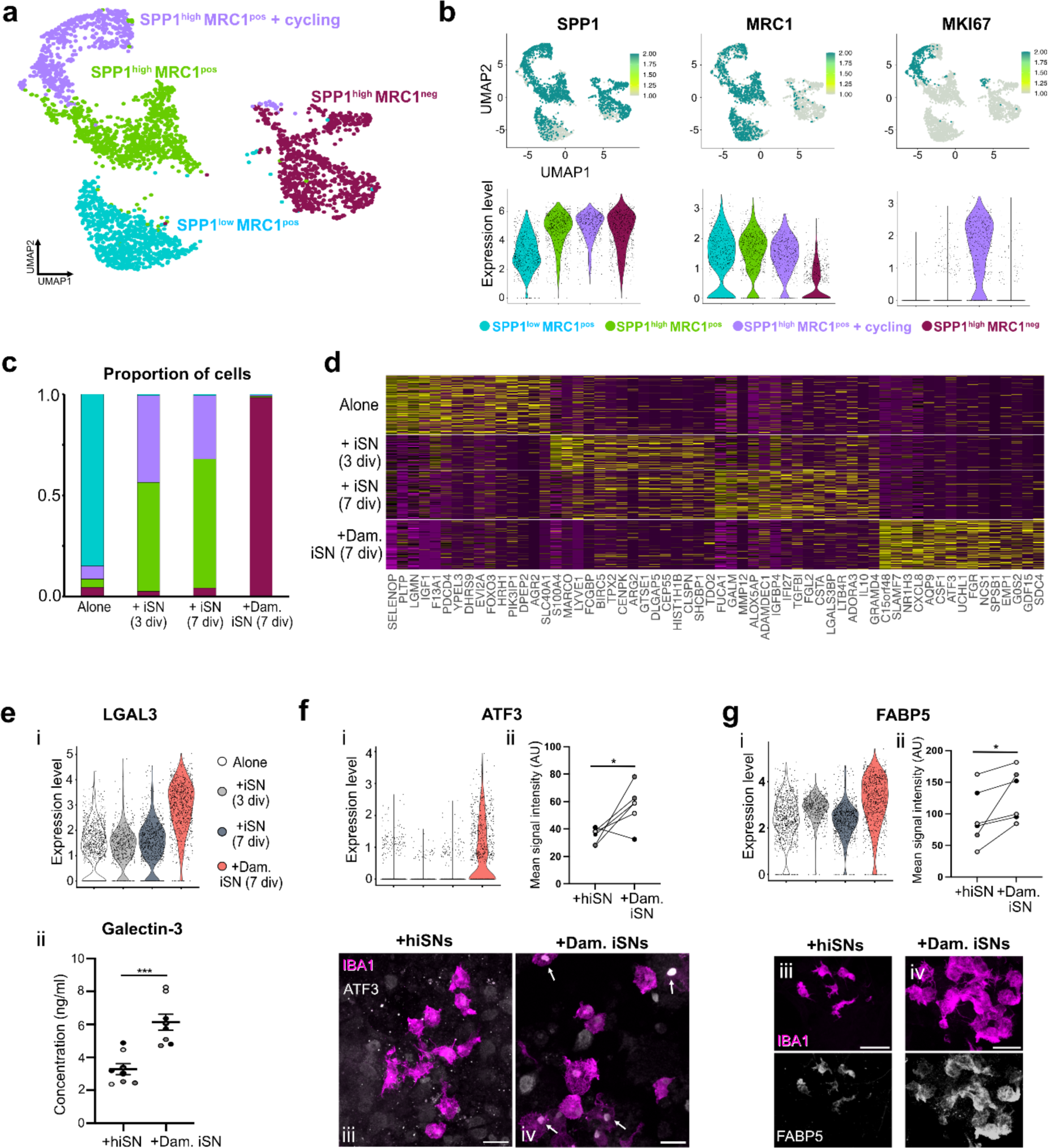
iMac gene expression profile is driven by co-culture condition. **a** UMAP plot of iMacs from all conditions showing four discrete clusters (n=1 differentiation and 1 iPSC line): total cells (3157), cultured alone (987 cells), with healthy iSNs for 3 (576 cells) and 7 days (791 cells), and for 7 days with injured iSNs (803 cells). **b** UMAP feature plots and corresponding violin plots of the genes used to describe each cluster. Each dot is a cell that was sequenced; green dots indicate those positive for a given gene. **c** Proportion of iMacs from different conditions assigned to each of the four clusters. Note that most iMacs from each condition were assigned to one cluster (or two in the case of co-culture with healthy iSNs). **d** Heatmap showing gene expression profile organised by iMac condition. The top fifteen differentially expressed genes per condition are shown. **e** (i) Violin plot and validatory immunoassay (ii) for Galectin-3, showing its upregulation in iMacs co-cultured with damaged iSNs. The immunoassay was performed on supernatant from div 7 co-cultures. Data represent mean ± SEM, with values for individual differentiations also shown (n=8, from 3 independent iPSC lines: *black circles - SBAD2, grey circles - SBAD3 and white circles - ZsGreen*). Two-tailed unpaired *t*-test. \*\*\**P*<0.05. **f** (i) Violin plot showing upregulation of the injury gene ATF3 in iMacs co-cultured with injured iSNs. Quantification of mean ATF3 signal intensity (ii) and representative images of iMacs co-cultured with healthy (iii) or injured iSNs (iv) for 7 days illustrating cells with nuclear ATF3 (white arrows). **g** (i) Violin plot for FABP5. Quantification of mean FABP5 signal intensity (ii) and representative images of iMacs co-cultured with healthy (iii) or injured iSNs (iv) for 7 days illustrating FABP5 expression. For quantification in **f** and **g**, paired values represent independent differentiations in which co-culture, ICC processing and image analysis were performed in parallel (n=6, from 3 independent iPSC lines: *black circles - SBAD2, grey circles - SBAD3 and white circles - ZsGreen*). Two-tailed paired *t*-test. \**P*<0.05. *Scale bars represent 25*µm.

Macrophages can be characterised as tissue-resident or infiltrating. Following peripheral nerve injury, the site of the lesion is characterised by extensive monocyte infiltration and proliferation of resident macrophage populations; moreover DRG-resident macrophages have also been shown to proliferate, albeit to a lesser extent^8,10,11,53^. While we used iMacs, which more closely resemble tissue-resident macrophages for our primary model, we sought to test whether monocyte-derived macrophages (MoMs) responded similarly to co-culture with iSNs. MoMs derived from a healthy female donor were successfully maintained in co-culture with iSNs (Supp. Fig. 5a). Similar to iMacs, MoMs altered their gene expression dependent on the presence and state of iSNs (Supp. Fig. 5b-e and Supplementary Data Table 2). While clusters were described by different molecular markers, many of the genes enriched in MoMs co-cultured with damaged iSNs were also found to be enriched in iMacs under similar conditions (Supp. Fig. 5f and Supplementary Data Table 3). ATF3 and LGAL3, but not FAPB5, were found enriched in the damaged neuron condition (Supp. Fig. 5g). Taken together, these data illustrate that iMac gene expression is highly plastic and dependent on neuronal context, with iMacs cultured with damaged iSNs taking on a putatively more pro-inflammatory profile, features of which are shared with MoMs.

### Validation with *in vivo* murine datasets

While there is evidence that macrophage populations are involved in pain progression across different types of nerve injury^6,54,55^, their role is likely context-dependent and may evolve during the transition from initiation to maintenance of neuropathic pain. To better understand whether our *in vitro* system recapitulates features of the *in vivo* scenario, we profiled macrophage populations in the sciatic nerve of mice following sham surgery, as well as at early (1 week) and late (15 weeks) timepoints following two types of mechanical nerve trauma: partial sciatic nerve ligation (PSNL), which results in a non-resolving injury, and nerve crush, which allows nerve regeneration and recovery. We collected live single myeloid cells based on CD45^+^/CD11b^+^ staining (excluding Ly6G^+^ neutrophils at acute timepoints) (Supplementary Figs. 6 and 7) from sciatic nerve of sham and nerve-injured animals. Cells from different conditions were labelled with hashtag oligos and combined as described in *Methods*. Consistent with our *in vitro* data describing naïve clusters with MRC1, Mrc1^+^ resident macrophage clusters, characterized by expression of Cd163, Lyve1 and MHCII were the predominant populations present following sham surgery. At the acute timepoint, PSNL was characterised by an enrichment of an Spp1^+^Lgals3^+^ cluster and the presence of putative phagocytic cells (pMφ), whereas induced MHCII macrophages (iMHCII^+^ Mφ) predominated in the crush, alongside a clear enrichment for Alox15^+^ putative wound-healing macrophages (likely reflecting the favourable regenerative conditions of the crush model) (Fig. 4a-c and Supp. Fig. 8a). With both injury states, we also detected an expansion of Spp1^+^ CD11c^+^ macrophages, which resemble injury-associated microglia described in the CNS^39^. Finally, we detected small numbers of putative dendritic cells (DC), non-myeloid cells that escaped our sorting strategy (Il2ra+) and Lyc6+ monocytes. The last cell type likely appears rarer than it is *in vivo*, because monocytes die preferentially during scRNA-seq workflows. At the chronic time point, the differences between injury states appear to subside, with resident populations once more predominating across most samples.

**Figure 4:**
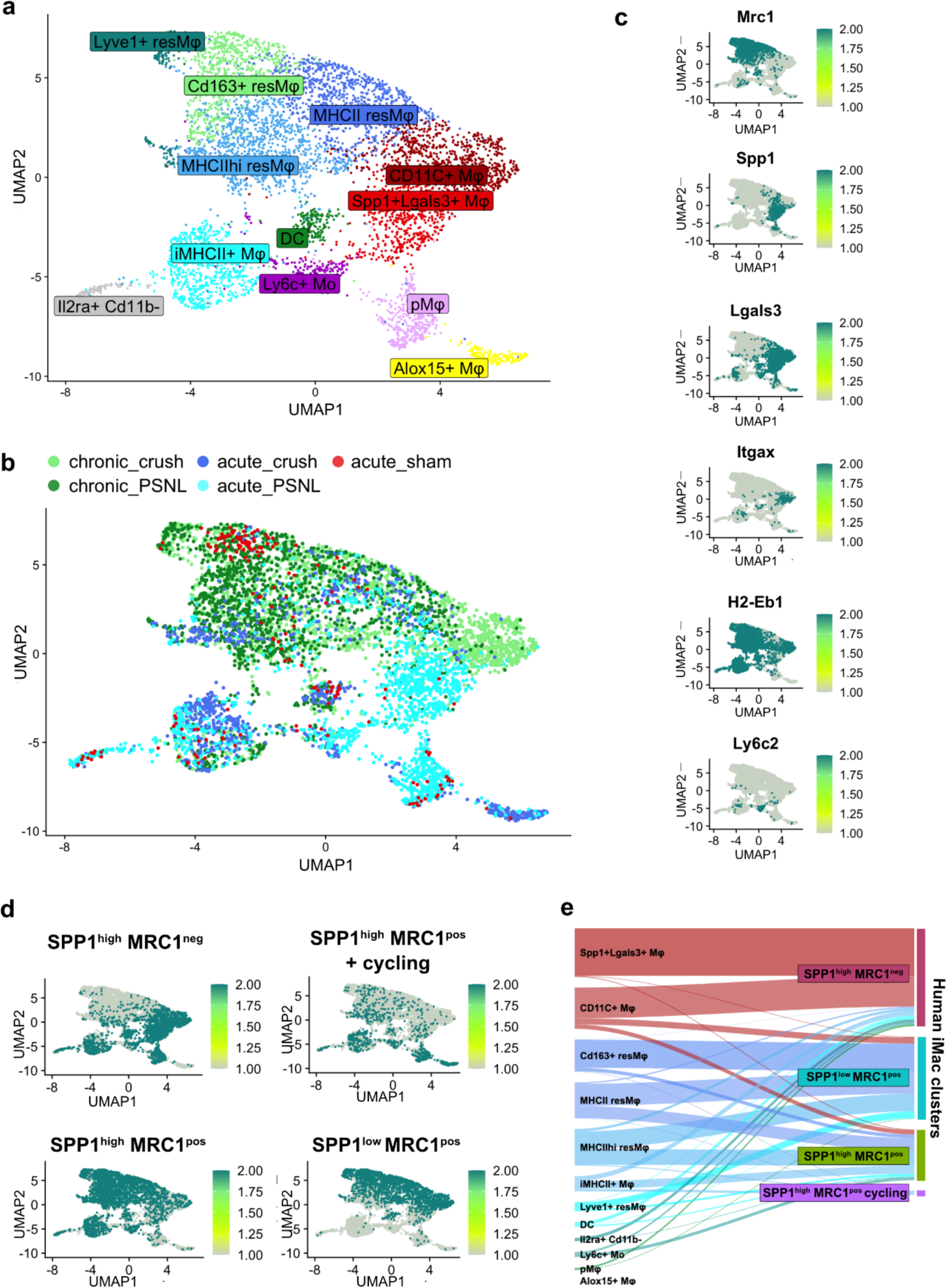
Mouse scRNA-seq data reveal that Spp1+ and Mrc1 macrophages are also a feature of nerve injury *in vivo*. ***a*** UMAP plot of CD11b+ myeloid cells derived from mouse nerve after injury at acute (1 week) and chronic (15 weeks) timepoints. **b** Injury state superimposed on the UMAP space. Each dot is a cell, the colour of each dot denotes which experimental group the cell derived from: 1 week (acute) post-sham, or 1 week (acute) or 15 weeks (chronic) post-PSNL or sciatic nerve crush injury. **c** Gene expression of several marker genes plotted onto the UMAP clusters. Each dot is a cell that was sequenced; green dots indicate those positive for a given gene. **d** Marker genes for the iMac clusters projected onto the mouse scRNA-seq UMAP space using the UCell package. To be included in the analysis, marker genes had to be expressed in at least 10% of cells in each cluster and had to differ in their expression from the other clusters by at least 20% points at adjusted *p* < 0.05. Injury-induced SPP1^high^MRC1^neg^ iMacs map well onto mouse injury-induced macrophages (majority blue clusters in b), while SPP1^low^MRC1^pos^ and SPP1^high^MRC1^pos^ map more prominently onto mouse resident (res) macrophage clusters labelled in a. **e** Sankey plot showing how mouse macrophage scRNA-seq clusters obtained from sciatic nerve after injury map onto the clusters observed in our human iMac clusters. Mouse macrophage clusters on the left, human macrophage clusters on the right. Of all 6,334 cells which were seeded from the human data, 65% could not be assigned and are not included here; it is unclear to what extent this is due to true biological differences or due to technical challenges associated with batch variability and orthologue conversion.

We sought to assess the similarity of our human macrophage populations to their mouse counterparts. Consistent with the human *in vitro* data, Spp1 expression was enriched in mouse injury-induced macrophage clusters, whereas, conversely, Mrc1 was enriched in resident macrophage populations that were abundant in the uninjured state (Fig. 4c). We projected marker genes from our human clusters onto the mouse scRNA-seq UMAP space using the UCell package. Injury-induced SPP1^high^MRC1^neg^ iMacs mapped well onto mouse clusters that were enriched at acute timepoints following PSNL and crush, while healthy iSN-associated SPP1^high^MRC1^pos^ iMacs mapped more prominently onto mouse naive-state resident macrophage clusters (Fig. 4d). Global expression profile alignment illustrated that injury-enriched mouse clusters mostly mapped to injury-induced SPP1^high^MRC1^neg^ iMacs, while resident macrophage populations map to both iMacs cultured alone (SPP1^low^MRC1^pos^) and iMacs cultured with healthy iSNs (SPP1^high^MRC1^pos^). Finally, we observed enrichment of our three neuropathic pain-associated genes (Lgals3, Atf3 and Fabp5) across murine injury states (Supp. Fig. 8b). These data illustrate the heterogeneity and dynamic nature of the macrophage response to nerve injury *in vivo*. They also demonstrate shared features between our injury-induced iMac populations and macrophage clusters observed at acute timepoints following nerve injury *in vivo,* supporting the validity of the *in vitro* model.

### iMacs augment hyperexcitability associated with neuron damage

Hyperexcitability and ectopic activity of sensory neurons are fundamental drivers of neuropathic pain^4,56,57^. Macrophages secrete many soluble mediators that can sensitise sensory neurons but there is no causal evidence demonstrating that macrophages directly influence sensory neuron activity following damage. We performed patch clamp recordings to assess neuron injury-induced changes in excitability and whether the presence of iMacs further influenced neuronal activity (Fig. 5a). In terms of passive membrane properties, damaged iSNs exhibited a depolarised resting membrane potential, an increase in input resistance and a reduced capacitance (Fig. 5b-d). The latter likely reflects the loss of neurites and a shrinkage of soma size. Evoked action potential threshold (rheobase) was also reduced following neuronal damage (Fig. 5e and f); however, this result should be considered in the context of a reduced capacitance of these neurons. While these data provide evidence for hyperexcitability following damage, there were no differences in passive properties or evoked firing between injured iSN with or without iMacs (Fig. 5b-f). Uninjured nociceptors are usually electrically dormant in the absence of stimulation and even following nerve injury paradigms, spontaneous activity is only seen at low levels *in vitro*^56^, which contrasts with *in vivo* recordings^58^. *Ex vivo* mouse DRG neurons exhibiting spontaneous activity have greater input resistance than those that do not^59^ and so we asked whether our injury paradigm induced changes in spontaneous activity. A greater proportion of iSNs exhibited spontaneous activity following injury (3 of 42 healthy iSNs *vs* 19 of 87 damaged iSNs, *p*=0.046, Fisher’s exact test, with and without iMac groups combined), but there was no difference between conditions with or without iMacs (Fig. 5g). However, while incidence was no different, we observed a stark difference in the frequency of firing of those neurons exhibiting spontaneous activity. Damaged iSNs alone fired spontaneous action potentials (APs) at low frequency (6.8 ± 5.0 APs/min), whereas equivalent neurons co-cultured with iMacs exhibited firing at considerably increased frequency (40.5 ± 13.1 APs/min) (Fig. 5h-i). Together, these data suggest that neuronal damage induces hyperexcitability. They also show that spontaneous activity, a cardinal feature of neuropathic pain, can be directly augmented by macrophages.

**Figure 5.**
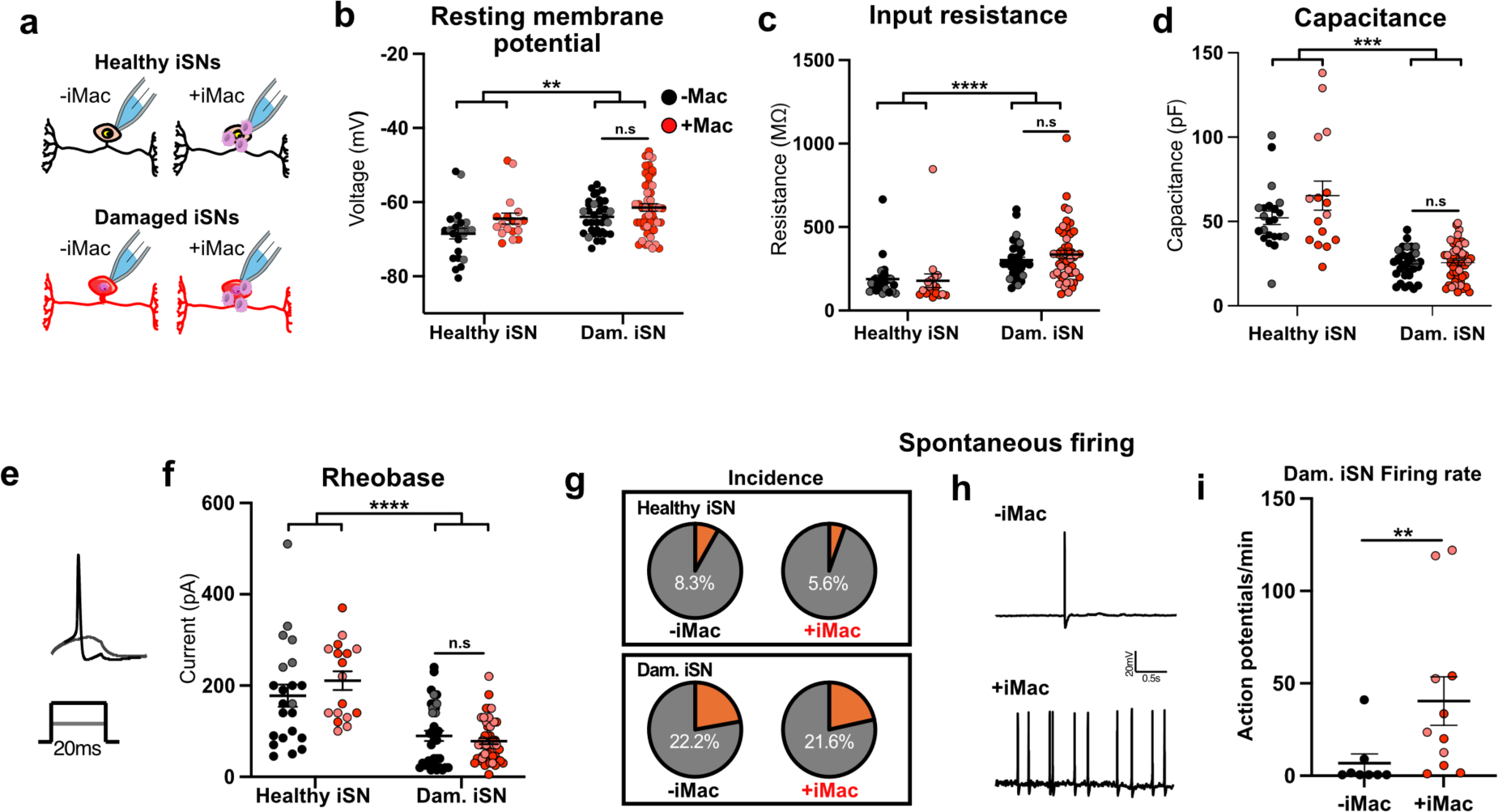
iMacs augment iSN spontaneous activity following injury. **a** Patch-clamp recordings were taken from healthy iSNs (hiSNs) and injured iSNs 7 days following injury, with or without iMac co-culture. **b, c, d** Passive membrane properties of iSNs across the conditions. Significance is shown for a two-way ANOVA analysis testing for main effects of injury and iMac presence and subsequent Tukey’s post-hoc tests (full numerical details of all tests are provided in Supplementary Table 1). **e** *R*epresentative image showing action potential firing of a healthy iSN to incremental current injections. **f** Mean ± SEM rheobase values. **g** Proportion of spontaneously active iSNs as defined by the firing of at least one action potential during a two-minute period. **h** Example traces of spontaneously active damaged iSNs without (*top*) and with (*bottom*) iMacs. **i** Action potential firing frequency of spontaneously active damaged iSNs. Mann-Whitney test. Note, healthy iSN groups could not be included in this analysis owing to samples only including 1-2 spontaneously active neurons. All data represent mean ± SEM with values for individual iSNs also shown (derived from 2 independent iPSC lines: *Black/red circles SBAD2, grey/light red circles-SBAD3*). \*\**P*<0.01, \*\*\**P*<0.001, \*\*\*\**P*<0.0001.

To look for evidence of intercellular communication, we performed ligand-receptor analysis with a recently published dataset of healthy iSNs at different stages of maturity^25^. Damaged iSN-associated SPP1^High^MRC1^neg^ iMacs had increased interactions compared to healthy iSN-associated SPP1^High^MRC1^pos^ iMacs, particularly for pathways associated with extracellular matrix signalling and cell adhesion (Supp. Fig. 9a-b). SPP1^High^MRC1^neg^ iMacs had higher interaction scores for several receptors with known capacity to directly sensitise sensory neuron activity, including CD44, ALK, ITGB1/TGAA3 and NRP1^60–63^ (Supp. Fig. 9b-c). CXCL8 (IL-8) has also been shown to heighten the excitability of sensory neurons^64^, although which receptor this requires is not yet known. These data lay the groundwork for future investigations of candidate signalling pathways involved in iMac influence on iSN excitability.

## Discussion

Here we describe the first humanised model to study the interactions between sensory neurons and macrophages; we use this to reveal a hitherto postulated but never empirically demonstrated ability of macrophages to directly sensitise the electrical activity of damaged sensory neurons. Injury to human (*in vitro*) and mouse (*in vivo*) sensory neurons initiates a major switch in associated macrophage identity, with overlap in gene expression across the systems validating our *in vitro* model. These data demonstrate the opportunity to use co-cultures of iSNs and iMacs for discovery science and as the basis for future work developing this system into an analgesic testing platform, usable at scale.

### Validity of the human co-culture model

Our results illustrating the plasticity of iMacs *in vitro* mirrors what has been observed by others when co-culturing them with CNS neurons^18,31,43^ and the remarkable extent to which macrophage identity *in vivo* is dependent on the local tissue environment^65^.

The SPP1^High^MRC1^neg^ macrophage profile abundant in co-cultures with damaged iSNs generally mapped well onto mouse clusters that are enriched in the damaged sciatic nerve at early timepoints following injury *in vivo*. SPP1 is not a typical classifier of peripheral nerve-associated macrophages but is a common marker of microglia associated with brain injury and disease^39,66^. We found Spp1 to be abundant in clusters enriched in our mouse samples of acute injury, consistent with results from previous studies^9^. One aspect where our human and mouse cell types differed was the expression of SPP1 in iMacs co-cultured with uninjured neurons. In mouse, Spp1 is not only associated with injury, but is also expressed by microglia close to emergent axon tracts during brain development^39^. iSNs exhibit features of a fully differentiated nociceptor at the time of our assays but are still maturing and in a period of neurite growth, providing a plausible explanation for why SPP1 expression was enriched in both conditions in our human system.

We found consistent up-regulation of neuropathic pain-associated genes (ATF3, LGALS3 and FABP5) across *in vitro* and *in vivo* experimental platforms of nerve injury, although FABP5 was not enriched in MoMs. Study of ATF3 following nerve injury has largely focussed on *de novo* neuronal expression, given the protein’s role in nerve regeneration^67,68^. However, ATF3 has a well described role in macrophage biology where it is thought to act as a repressor of pro-inflammatory gene expression and cytokine release^69,70^. LGALS3 encodes the multifunctional protein, Galectin-3, which is a TREM2 ligand with described roles in several aspects of macrophage biology. It has been implicated in various neurological diseases, including neuropathic pain, where knockout or pharmacological inhibition of Galectin-3 reduces mouse pain behaviours following nerve injury^48,71^. Beyond gene expression changes, we observed increased secretion of CCL3 and CCL4 from iMacs co-cultured with damaged iSNs. While the role of CCL2 as the primary monocyte/macrophage recruitment signal following peripheral nerve damage is well studied^72,73^, the contribution of CCL3 and CCL4 is less so. Knockout of CCL3 or the major cognate receptor of CCL4 (CCR5) has no effect on macrophage recruitment to damaged nerves^73,74^, yet signalling from both contributes to pain behaviours^37,38^. It is therefore possible that these chemokines function to modify macrophage profile^75,76^, or to directly sensitise sensory neurons^77,78^. All of these factors have previously been proposed as viable therapeutic targets^71,79,80^, and therefore our model system offers an attractive platform in which to test their contribution.

Few (if any) studies have profiled macrophage biology *in vivo* at chronic timepoints following different types of nerve injury. We observed that differences in macrophage clusters between resolving and persistent nerve damage appeared to subside over time, highlighting the evolving macrophage response. It is not clear how this relates to symptoms, with evidence suggesting that pain is not only limited to non-resolving nerve injuries, like PSNL, but can also arise from nerve crush^81^, as well as during re-innervation^82^. Mechanistically, it remains to be determined if and how the differences we observed in acute macrophage phenotypes influence other inflammatory and stromal cell types long-term.

### Macrophages directly amplify damaged sensory neuron ectopic activity

Sensory neurons undergo major transcriptional changes in response to injury *in vivo*^83,84^. These result in a regenerative phenotype characterised by hyperexcitability and *de novo* ectopic activity^84,85^. The latter is considered a key driver of pain in people^4^. Our *in vitro* mechanical trauma model similarly induced expression of canonical injury-associated genes, rapid neurite regeneration and hyperexcitability. The finding that axotomy (in the absence of iMacs) induced a low level of spontaneous activity is consistent with what has been observed in *ex vivo* recordings of rodent sensory neurons from experimental nerve injury models^86^ and human DRG neurons derived from individuals with neuropathic pain^87^ (both situations with limited macrophage presence), suggesting that at least some of the disease features of a damaged sensory neuron are cell intrinsic.

Macrophages have been proposed as a source of pro-inflammatory mediators following nerve injury^72,88^, some of which have been shown to sensitise sensory neurons in isolation^89–91^. However, we believe that our findings are the first to demonstrate the capacity of macrophages to directly amplify damaged sensory neuron hyperexcitability, in mouse or human. The finding that iMacs did not impact passive membrane properties or evoked firing properties, but did increase spontaneous (i.e. non-evoked) firing frequency could have implications for our understanding of the cell type’s role in nerve injury-induced pain.

Experimental depletion of macrophage populations reduces evoked hypersensitivities in preclinical models of nerve injury^6^, but behavioural readouts of spontaneous pain have not been examined. Both stimulus-evoked and paroxysmal pain scores of Morton’s neuroma patients correlate with a greater presence of macrophage populations in nerve tissue^7^. Spontaneous activity of sensory neurons is often linked to spontaneous pain^4^, and so considering our data, and that of Sandy-Hindmarch *et al*^7^, it may be prudent to re-evaluate the preclinical models to assess the role of macrophages in maintaining spontaneous pain. If our hypothesis is correct, then it would highlight disrupting pathological macrophage-sensory neuron signalling as an attractive approach to abrogate a major clinical complaint of people living with neuropathic pain^92^.

### Future opportunities

We limited our study to a mechanical trauma model, which aligns with preclinical work in which the most studied neuropathic models are similarly based on mechanical nerve damage. However, macrophage involvement has been postulated in systemic neuropathies, such as chemotherapy and diabetes-induced painful neuropathies^54,93^. Efforts have been made to model these insults *in vitro*^94–96^ and so it would therefore be of interest to test co-culture of iMacs and iSNs under these conditions.

Our model offers a new platform to study sensory neuron-macrophage cross talk, which is known to be important for a myriad of biological processes beyond pathological pain, including normal mechano- and thermo-transduction^97^, wound repair and fibrosis^98,99^, and defence against infection^100,101^. In nerve regeneration, macrophages are classically considered pro-regenerative due to their actions of debris clearance and signalling to Schwann cells^12,35,102,103^. A study by Hakim and colleagues has recently described a role for nerve-associated macrophages in protecting against sensory axon loss in a mouse model of diabetic peripheral neuropathy^93^. Of interest, this study identified Galectin-3 up-regulation as a key mediator of this role^93^. iMacs in our co-culture were highly efficient at clearing neuronal debris following damage and neurite regeneration appeared qualitatively changed, with neurites aligning into dense tracts. While we did not specifically address the contribution of iMacs to neuroprotection and neurite regeneration, this would be a fruitful avenue of investigation.

In summary, we present a humanised co-culture model to study the interactions of sensory neurons and macrophages. We have used this model to show a direct action of macrophages in sensitising the electrical activity of damaged sensory neurons and to identify candidate signalling pathways that may be amenable to therapeutic disruption.

## Acknowledgements

We thank John Craig for technical assistance performing immunoassays and Professor Lesley Forrester for gifting the SFCi55ZsG iPSC line. Schematics included in Figures 1, 2, 5 and Supp Fig. 2 include images that were adapted from images provided by Servier Medical Art, licensed under CC BY 4.0. The research was funded by an MRC Fellowship grant (MR/T01072X/1) and an Ono Pharmaceuticals Rising Stars Award held by GAW. PC was supported by University of Glasgow LKAS Leadership funds. ZH received support from an MRC New Investigator Award (MR/P010814/01) awarded to FD. JVW is funded by a Wellcome Trust Collaborative Award (224257/Z/21/Z). DS and MKS were funded by Versus Arthritis UK award (23229), KD and AE were funded by the Versus Arthritis UK award (22072). For the purpose of Open Access, the author has applied a CC BY public copyright licence to any Author Accepted Manuscript (AAM) version arising from this submission.

## Supplementary figures

**Supplementary figure 1.**
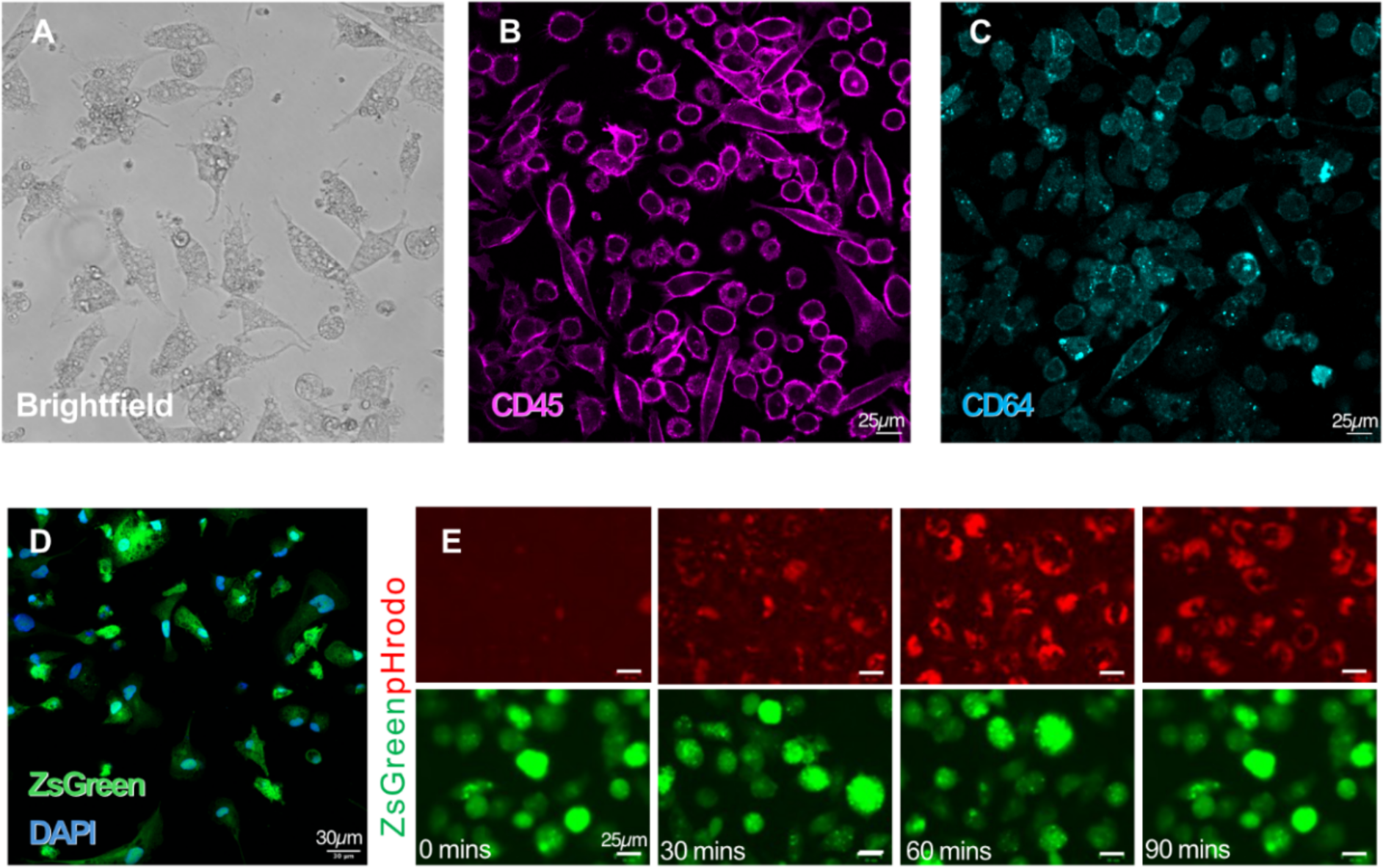
iMac generation. **a-c** Representative brightfield and fluorescent images of macrophage-like cells (iMacs). Cells stain positive for the hematopoietic cell markers CD45 and CD64. **d** Representative image of iMacs produced from the ZsGreen knock-in line. **e** Time lapse of ZsGreen-derived iMacs incubated with pHrodo Red E. coli BioParticles, which fluoresce in acidic environments (such as lysosomes). The gradual increase in pHrodo signal indicates iMacs are competent phagocytes.

**Supplementary figure 2.**
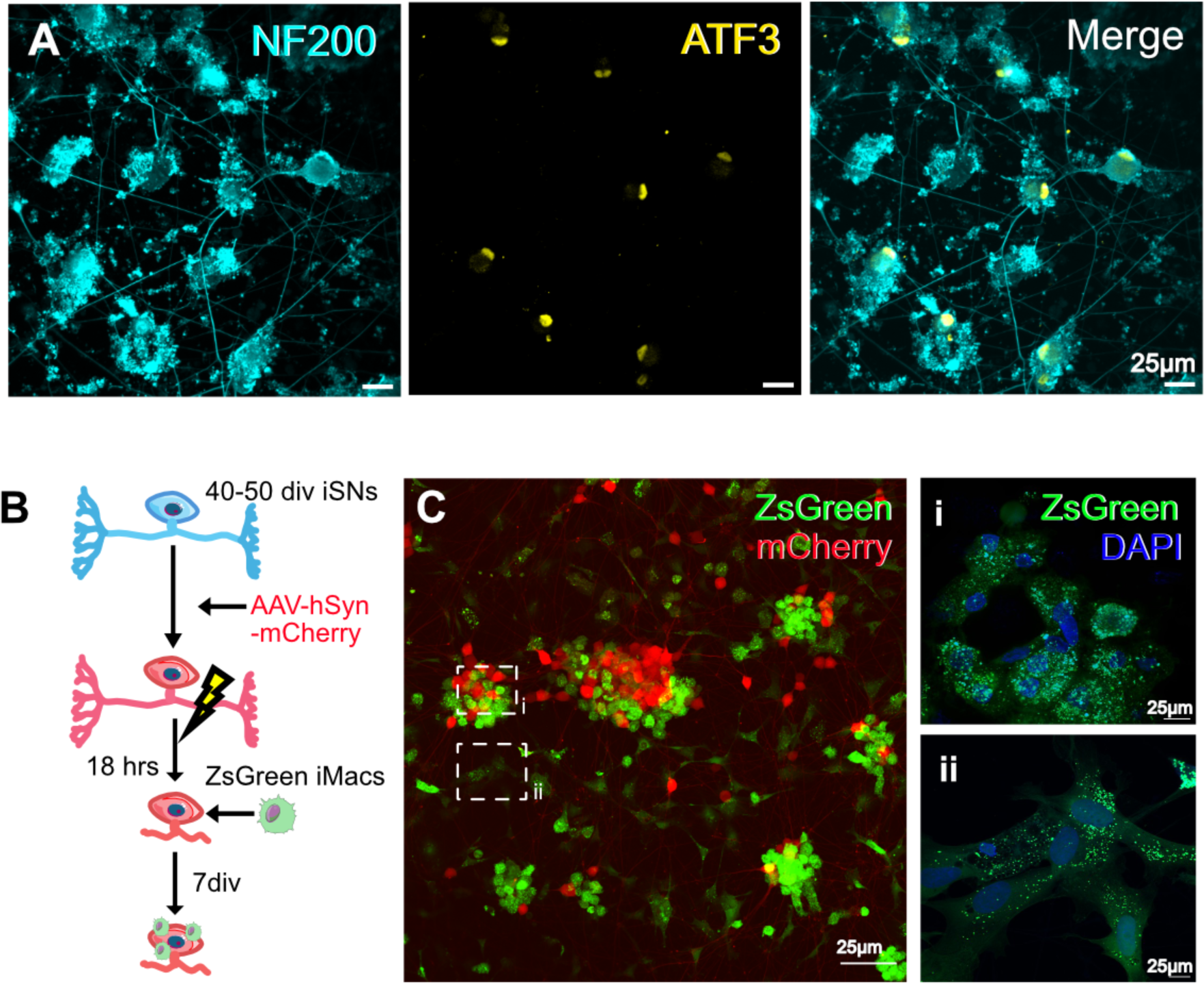
Co-culture of iMacs with damaged iSNs. **a** iSNs 24hrs after axotomy and replating, illustrating expression of the canonical injury marker, ATF3, and evidence of rapid neurite regeneration. **b** Schematic of mechanical trauma model in which neurons have been transduced with AAV-hSyn-mCherry prior to axotomy and replating, and subsequent co-culture with ZsGreen iMacs. **c** Representative image of 7 div co-culture. Most iMacs aggregate around neuron soma clusters and appear to exhibit an ameboid profile (**i**), with iMacs not associated with these clusters exhibiting a larger, flattened morphology (**ii**).

**Supplementary figure 3.**
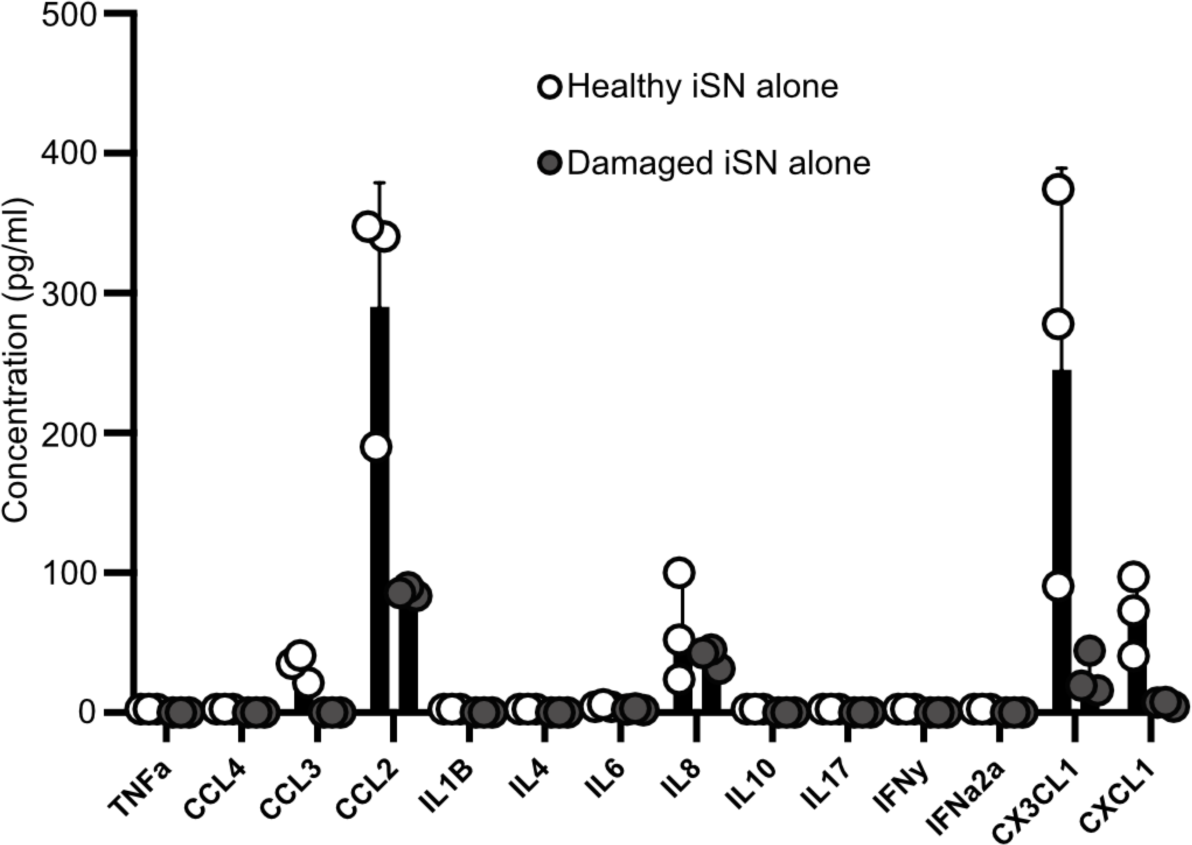
iSN monoculture secretory profiles. Discovery multiplex immunoassay detecting pain-relevant cytokines/chemokines in the supernatant after 3 days of monoculture in neuronal media + M-CSF. Data represent mean ± SEM (n=3, pooled from SBAD2 and SBAD3 iPSC lines). Relates to Fig 2 **e** but note the different Y-axis scale.

**Supplementary figure 4.**
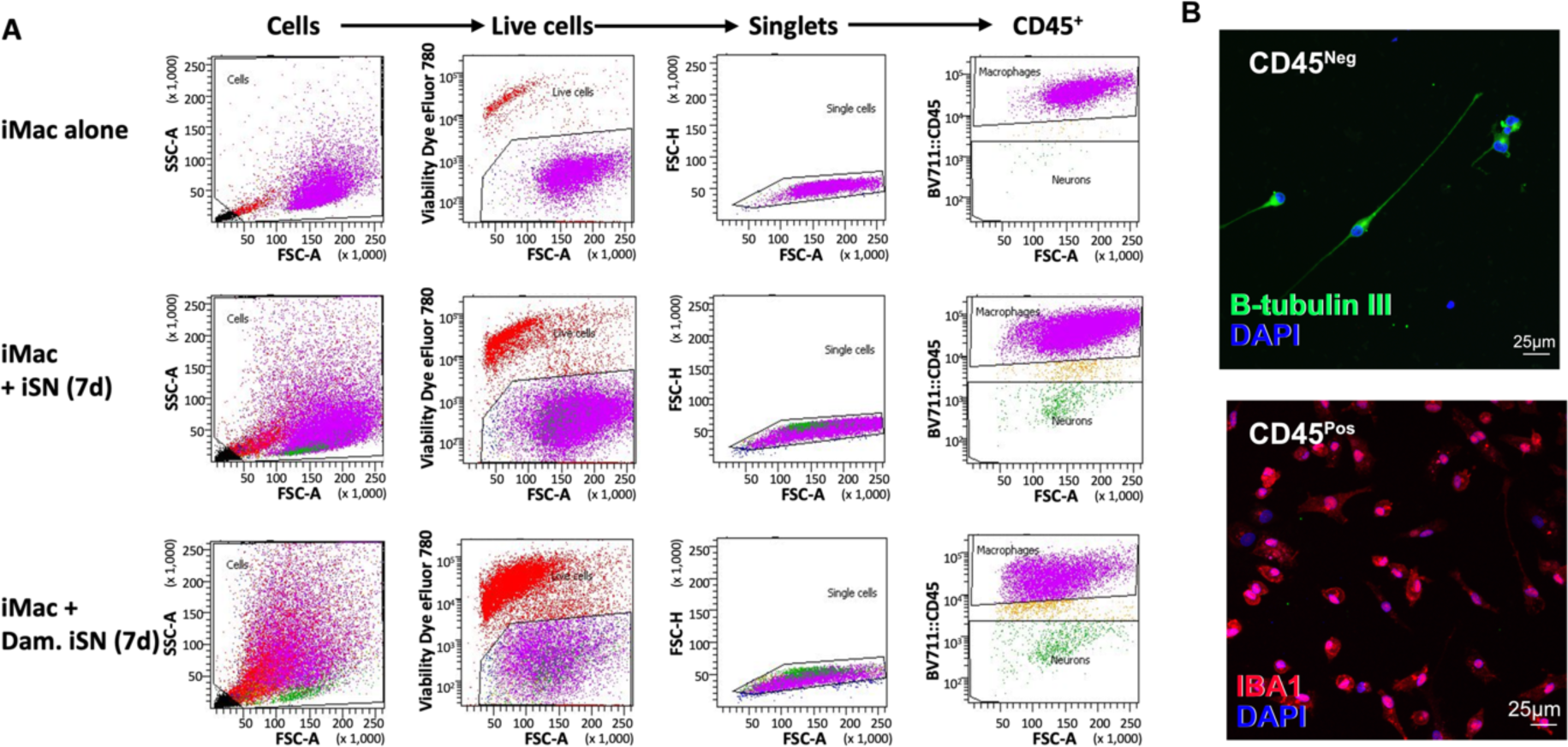
FACS gating strategy for iMac samples collected for scRNA-seq. **a** Samples containing iMacs cultured alone or with healthy or damaged iSNs for 7 days were gated based on forward (FSC) and side scatter (SSC) to enrich events for cells, live cells (i.e. those which were Viability eFluor™ 780 negative), singlets, and CD45 positive events. Neurons were not routinely collected. **b** Representative image of cells taken from CD45 negative and positive FACS gates during a pilot experiment, confirming appropriate capture of the two cell types.

**Supplementary figure 5.**
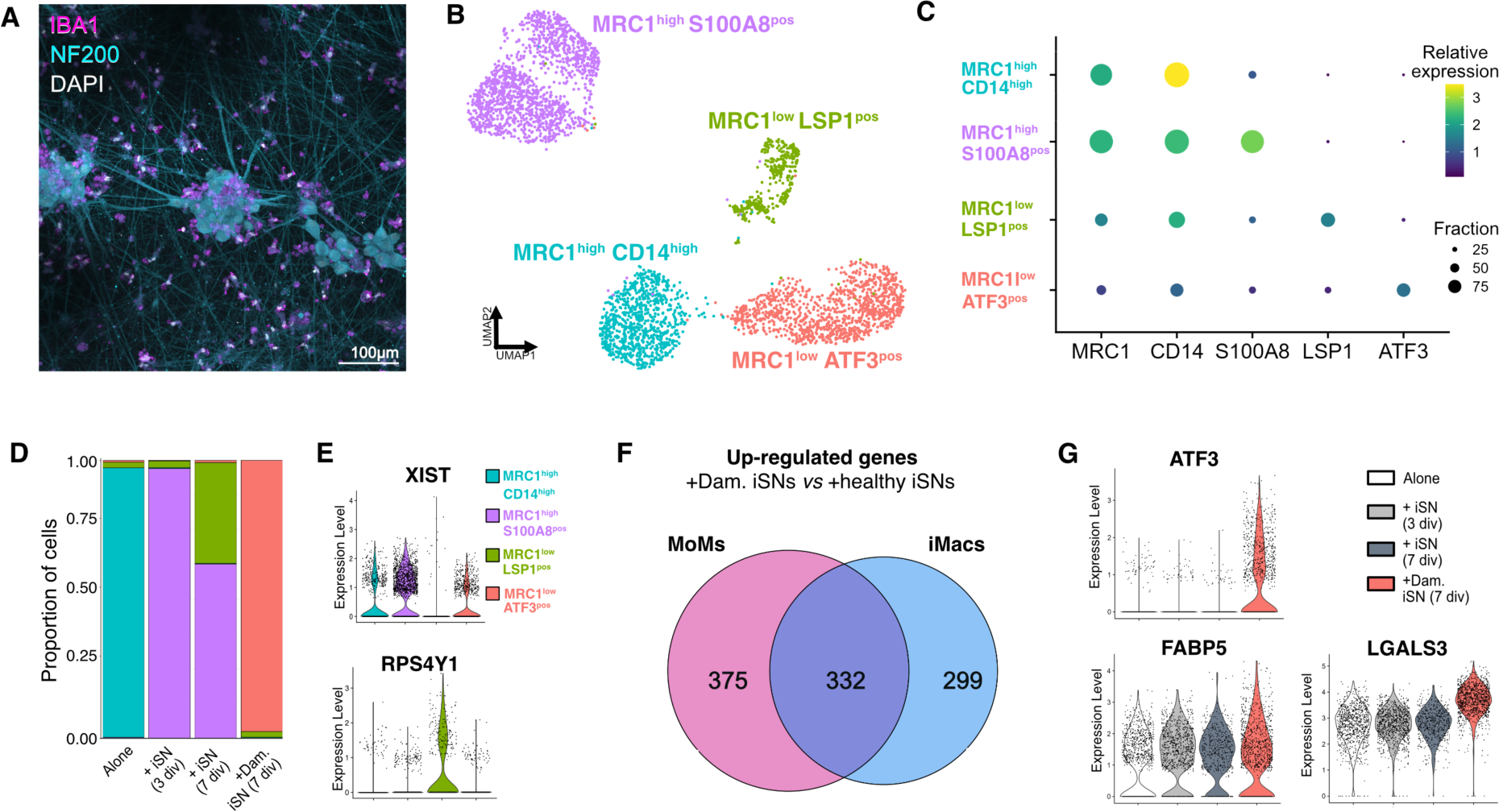
scRNA-seq analysis of MoMs. **a** Representative image of monocyte-derived macrophages (MoMs) co-cultured with iSNs for 7 days under conditions identical to iMac co-culture. MoMs stain positive for the macrophage marker IBA1. **b** UMAP projection of MoMs pooled from 4 conditions; cultured alone (895 cells) and co-cultured with healthy iSNs for 3 (980 cells) and 7 (854 cells) days, or with injured iSNs for 7 days (1113 cells) (Note, n=1 differentiation and 1 donor). Four distinct clusters can be seen. **c** Dot plot showing the relative expression of markers used to define clusters. **d** Proportion of MoMs from different conditions corresponding to each of the four clusters. Most clusters consist of MoMs from one condition, similarly to iMacs. **e** We noted that unlike other clusters, the MRC1^Low^LSP1^pos^ cluster had low expression of the female-biased gene XIST and enriched expression of the Y-linked gene RPS4Y1, leading to a reasonable suspicion that the female donor derived sample was partially (<10% cells) contaminated with male donor-derived MOMs concurrently in culture. We therefore excluded the MRC1^Low^LSP1^pos^ cluster from subsequent analysis. **f** Venn diagram of genes upregulated in either iMacs or MoMs when co-cultured with healthy *vs* injured iSNs (FDR < 0.05 and log_2_FC > 0.5), showing an overlap of approximately 50%. This is far greater than expected by chance: hypergeometric *p* value = 3.289669e-238. **g** Violin plots of three injury-associated genes found to be upregulated in iMacs co-cultured with injured iSNs. ATF3 and LGALS3 are also upregulated in MoMs cultured with damaged iSNs, however no difference is observed in FABP5 expression.

**Supplementary figure 6.**
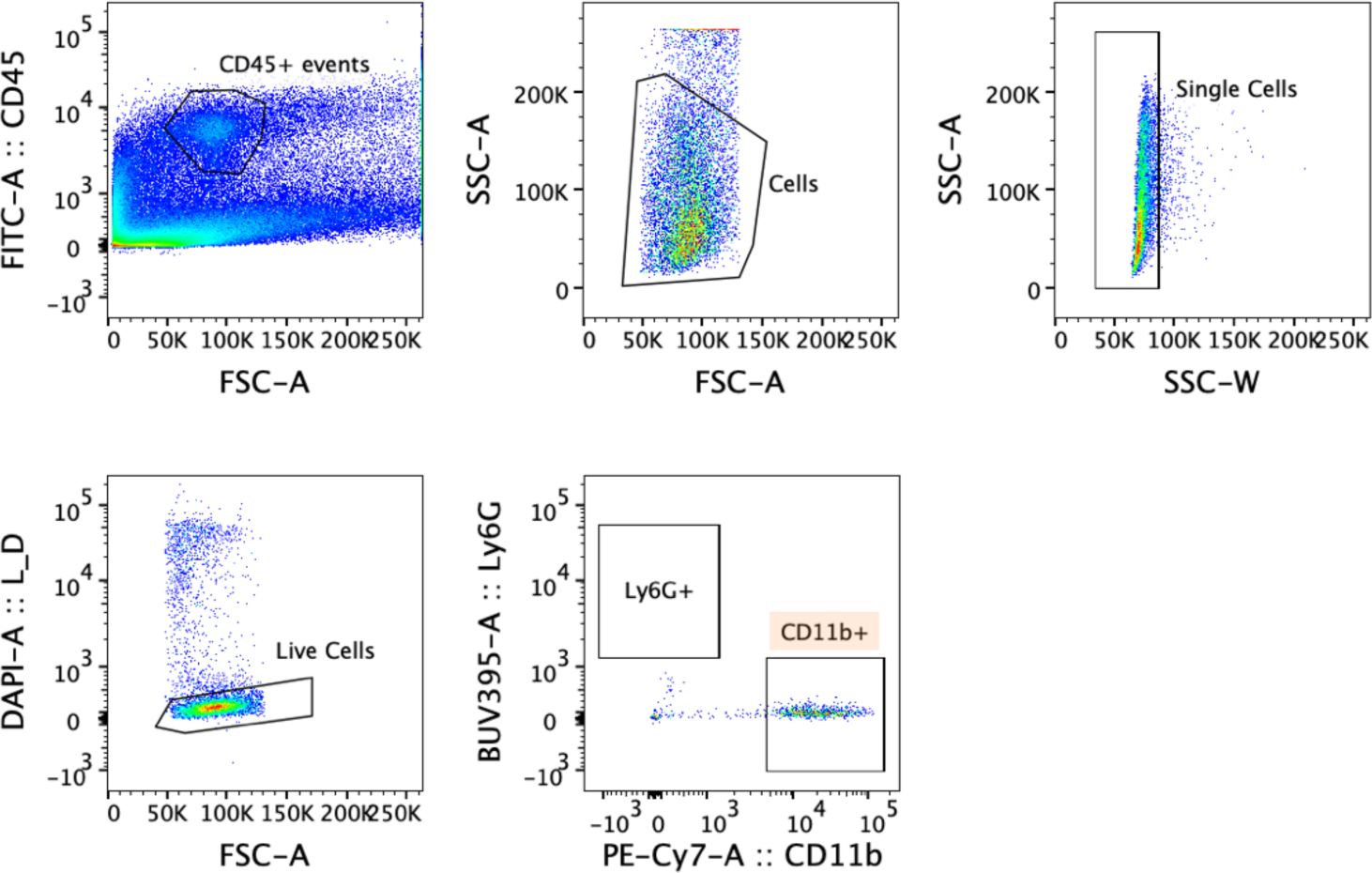
FACS gating strategy for samples taken at 1-week post-sciatic nerve injury. Data shown here derive from a single female PSNL mouse. CD45+ events were selected, followed by events that are likely cells based on forward (FSC) and side scatter (SSC), single cells based on the area (SSC-A) and width (SSC-W) of the side scatter, and live cells, i.e. those which were DAPI negative. After this, we sorted cells which were negative for the neutrophil marker Ly6G, but positive for the myeloid cell marker CD11b. All gates were placed on fluorescence minus one controls. At day 7 post nerve injury, we were not able to observe neutrophils, in keeping with what we have observed before in the PSNL model, where neutrophil infiltration appears to peak at day 1 post injury (Liang et al., Pain, 2020, 10.1097/j.pain.0000000000001914).

**Supplementary figure 7.**
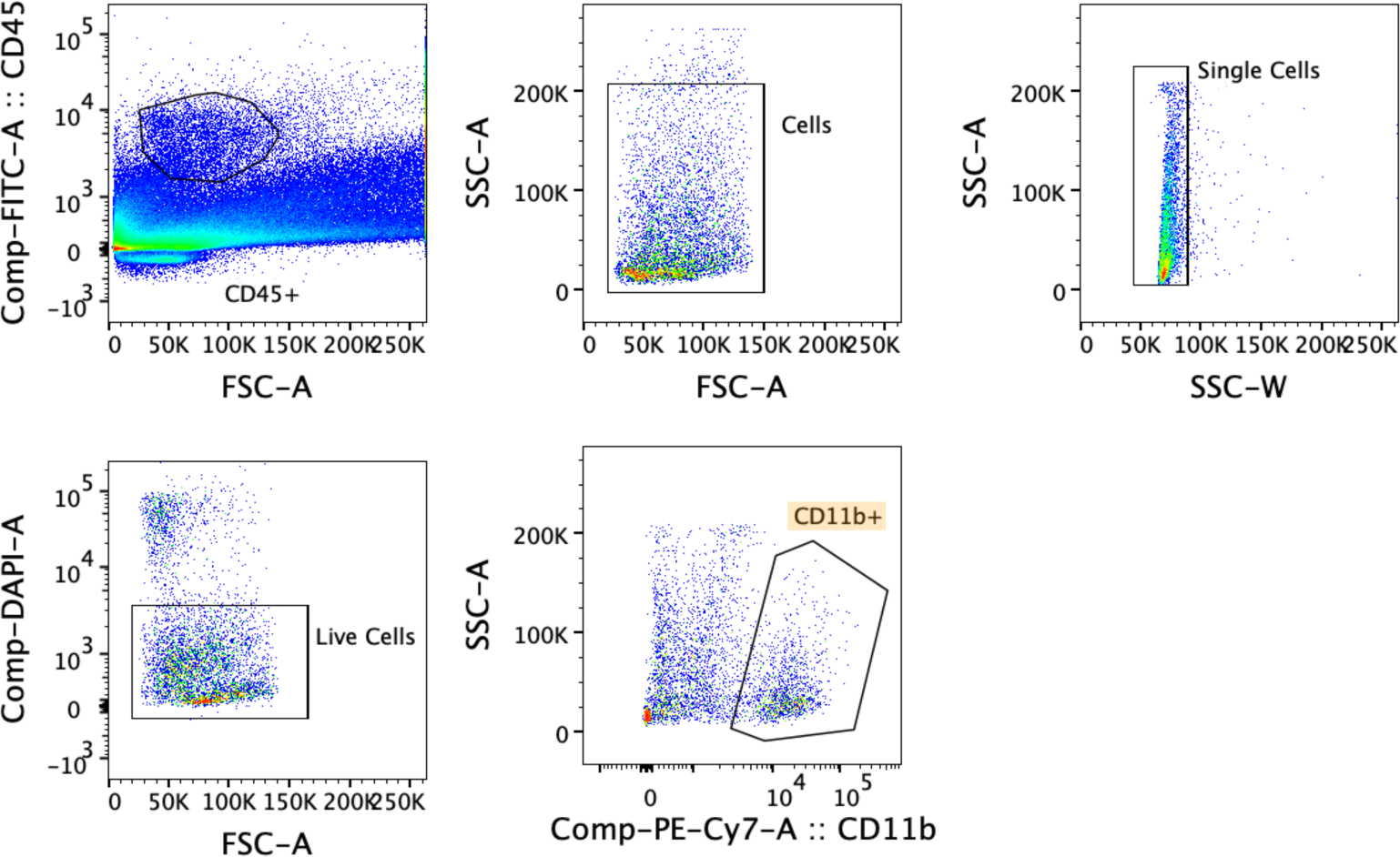
FACS gating strategy for samples taken at 15-weeks post-sciatic nerve injury. Data shown here derive from a single female PSNL mouse. CD45+ events were selected, followed by events that are likely cells based on forward (FSC) and side scatter (SSC), single cells based on the area (SSC-A) and width (SSC-W) of the side scatter, and live cells, i.e. those which were DAPI negative. After this, we sorted cells which were positive for the myeloid cell marker CD11b All gates were placed on fluorescence minus one controls.

**Supplementary figure 8:**
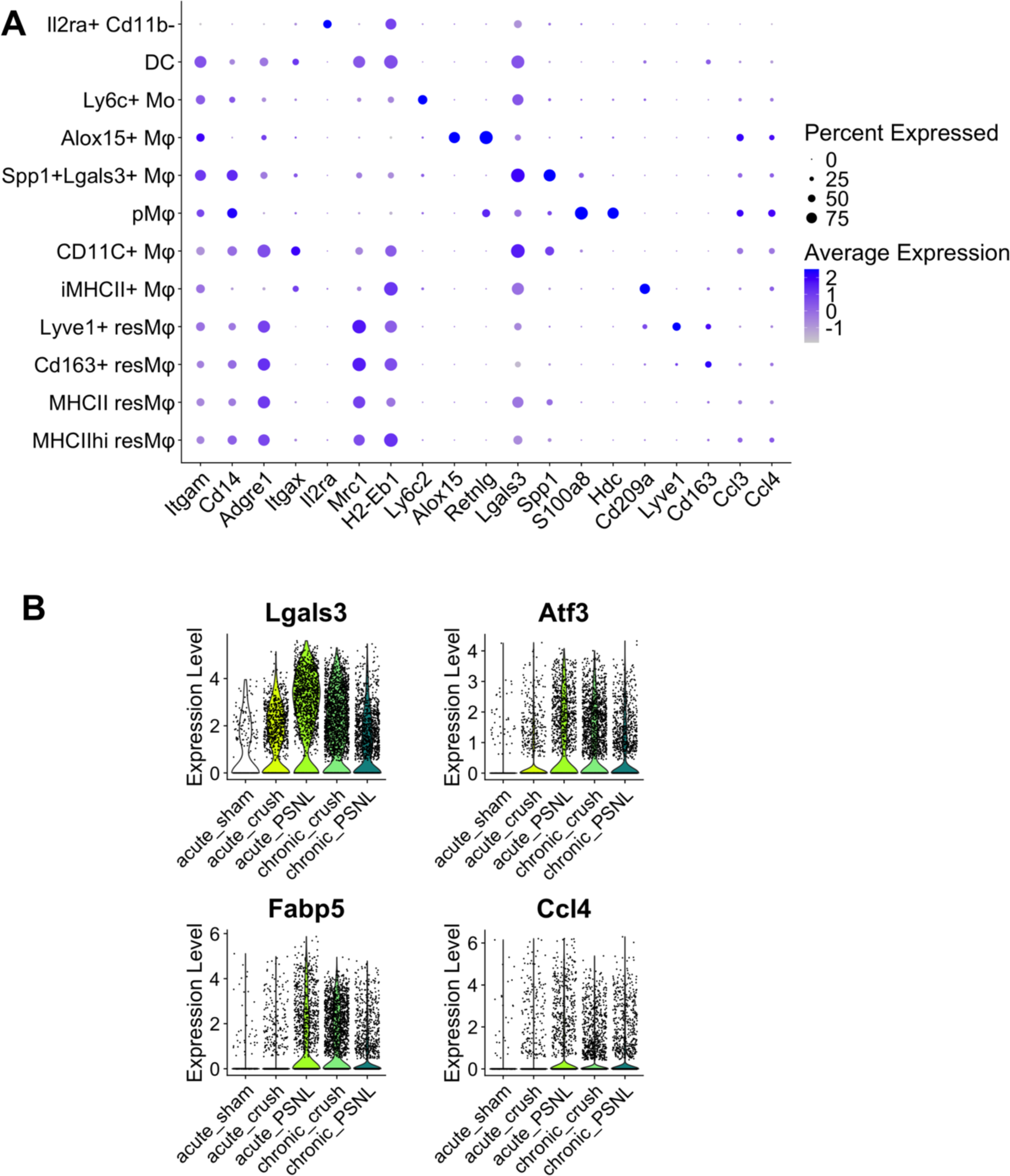
Mouse scRNA-seq data reveal that equivalent marker genes can be detected *in vivo* to those observed in our human iMac model. **a** Plots showing the expression level of marker genes in different cell clusters observed after sequencing mouse myeloid cells obtained from sciatic nerve either after sham surgery or 1-week or 15-weeks after nerve damage. Mφ = macrophages, p = phagocytic, i = injury-induced, Mo = monocytes, DC = dendritic cells and res = resident. **b** Violin plots of Lgals3, Atf3, Fabp5 and Ccl4 expression in the different injury conditions: samples taken one week after sham, crush or PSNL surgery (acute) vs. samples taken 15 weeks after crush or PSNL surgery (chronic).

**Supplementary Figure 9.**
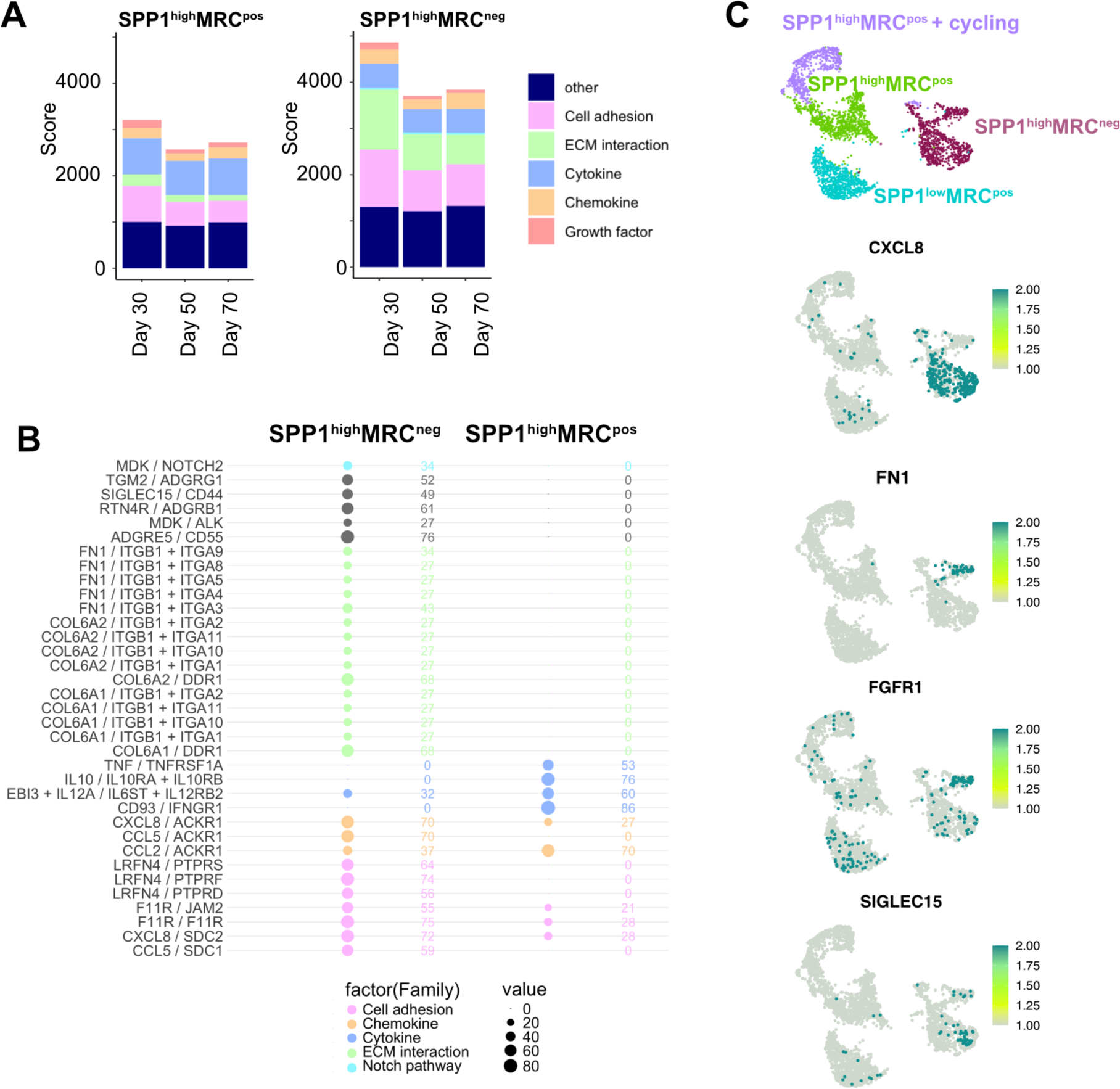
Ligand receptor interactions. **a** Communication scores for ligand-receptor interactions between iSNs differentiated for different lengths of time and SPP1^high^MRC^pos^ (which predominate in the healthy iSN condition) and SPP1^high^MRC^neg^ iMacs (which predominate in the damaged iSN condition). iSN RNA-seq data originally published by Li et al., Pain, 2025, 10.1097/j.pain.0000000000003512; day 50 corresponds to the age of the iSNs grown in our functional experiments. Scores are plotted for different ligand-receptor families with higher scores representing a greater number of matching ligand-receptor pairs expressed by the sender (iMac population) and receiver (iSNs). **b** Top differential ligand receptor pairings comparing the two iMac populations. The colours denote the function of each ligand-receptor pair, the size of the circles and corresponding numbers represent the communication scores. **c** UMAP feature plots of ligands with enriched expression in SPP1^high^MRC^neg^ for which there is prior data implicating either the ligand or a cognate receptor in sensory neuron excitability.

**Supplementary Table S1.** Two-way ANOVA degrees of freedom and *P* values for analyses of the effects of injury (healthy *vs* damaged iSNs) and/or presence of iMacs (related to Figure 5).

| Parameter | Injury | iMac presence | Interaction (Injury x iMac presence) |
| --- | --- | --- | --- |
| **Resting membrane potential** | $F_{1, 123} = 9.226$<br>$P=0.0029$ | $F_{1, 123} = 7.182$<br>$P=0.0084$ | $F_{1, 123} = 0.3720$<br>$P=0.5430$ |
| **Input resistance** | $F_{1, 121} = 22.78$<br>$P<0.0001$ | $F_{1, 121} = 0.1844$<br>$P=0.6684$ | $F_{1, 121} = 0.5834$<br>$P=0.4465$ |
| **Capacitance** | $F_{1, 115} = 97.94$<br>$P<0.0001$ | $F_{1, 115} = 4.143$<br>$P=0.0441$ | $F_{1, 115} = 3.354$<br>$P=0.0696$ |
| **Rheobase** | $F_{1, 119} = 58.45$<br>$P<0.0001$ | $F_{1, 119} = 0.5362$<br>$P=0.4654$ | $F_{1, 119} = 2.348$<br>$P=0.1281$ |

